# Noise schemas aid hearing in noise

**DOI:** 10.1101/2024.03.24.586482

**Authors:** Jarrod M. Hicks, Josh H. McDermott

## Abstract

Human hearing is robust to noise, but the basis of this robustness is poorly understood. Several lines of evidence are consistent with the idea that the auditory system adapts to sound components that are stable over time, potentially achieving noise robustness by suppressing noise-like signals. Yet background noise often provides behaviorally relevant information about the environment, and thus seems unlikely to be completely discarded by the auditory system. Motivated by this observation, we explored whether noise robustness might instead be mediated by internal models of noise structure that could facilitate the separation of background noise from other sounds. We found that detection, recognition, and localization in real-world background noise was better for foreground sounds positioned later in a noise excerpt, with performance improving over the initial second of exposure to a noise. These results are consistent with both adaptation-based and model-based accounts, since both explanations require online noise estimation that should benefit from acquiring more samples. However, performance was also robust to interruptions in the background noise and was enhanced for intermittently recurring backgrounds, neither of which would be expected from known forms of adaptation. Additionally, the performance benefit observed for foreground sounds occurring later within a noise excerpt was reduced for recurring noises, suggesting that a noise representation is built up during exposure to a new background noise and then maintained in memory. These findings suggest noise robustness is supported by internal models—“noise schemas”—that are rapidly estimated, stored over time, and used to estimate other concurrent sounds.

## Introduction

Much of the everyday listening experience is distorted by noise. Although noisy environments present a challenge for hearing, human listening abilities are remarkably robust to noise, enabling us to converse over the hum of a restaurant or recognize sounds on a windy day. However, the ability to hear in noise is vulnerable, declining with age (1, 2) and following hearing loss (1, 3). Understanding the basis of noise-robust hearing and its malfunction is thus a central goal of auditory research.

Noise robustness has been well documented in humans. For instance, speech intelligibility falls off gradually with signal-to-noise ratio (SNR), but remains high even when background noise has comparable power to a concurrent speech signal (4). Additionally, some types of sounds are easier to hear in noise than others (5), and hearing is more robust to some types of noise than others (6–10). Moreover, neural correlates to this robustness have been discovered along the ascending auditory pathway of multiple species (11–20). Yet, despite recent interest in the factors that enable and constrain hearing in noise, the problem is not well understood in computational terms.

A common view is that the auditory system filters out or suppresses noise in order to recognize sources of interest. One possibility is that the brain has internalized typical properties of noise and, by default, suppresses them relative to the properties of other sound sources (9, 12, 17). For instance, because noise is often approximately stationary (i.e., being defined by statistical properties that are relatively constant over time), the auditory system could preferentially suppress stationary sounds, which might enable more robust recognition of other sounds in noise. We refer to this hypothesis as “fixed noise suppression”, the idea being that there are fixed filters that attenuate noise-like sounds (Fig. 1A). Another possibility is that noise properties are implicitly detected and suppressed via local adaptation mechanisms that reduce the response to features that are relatively constant in the auditory input (13, 14, 16, 18, 21–23). We refer to this hypothesis as “adaptive noise suppression” (Fig. 1B). While adaptive mechanisms can account for some of the observed neural responses to stimuli in noise, such proposals have primarily been evaluated with simple synthetic noise signals, leaving it unclear whether they explain robustness to noise sources containing the rich statistical structure present in natural environments (24).

**Figure 1.**
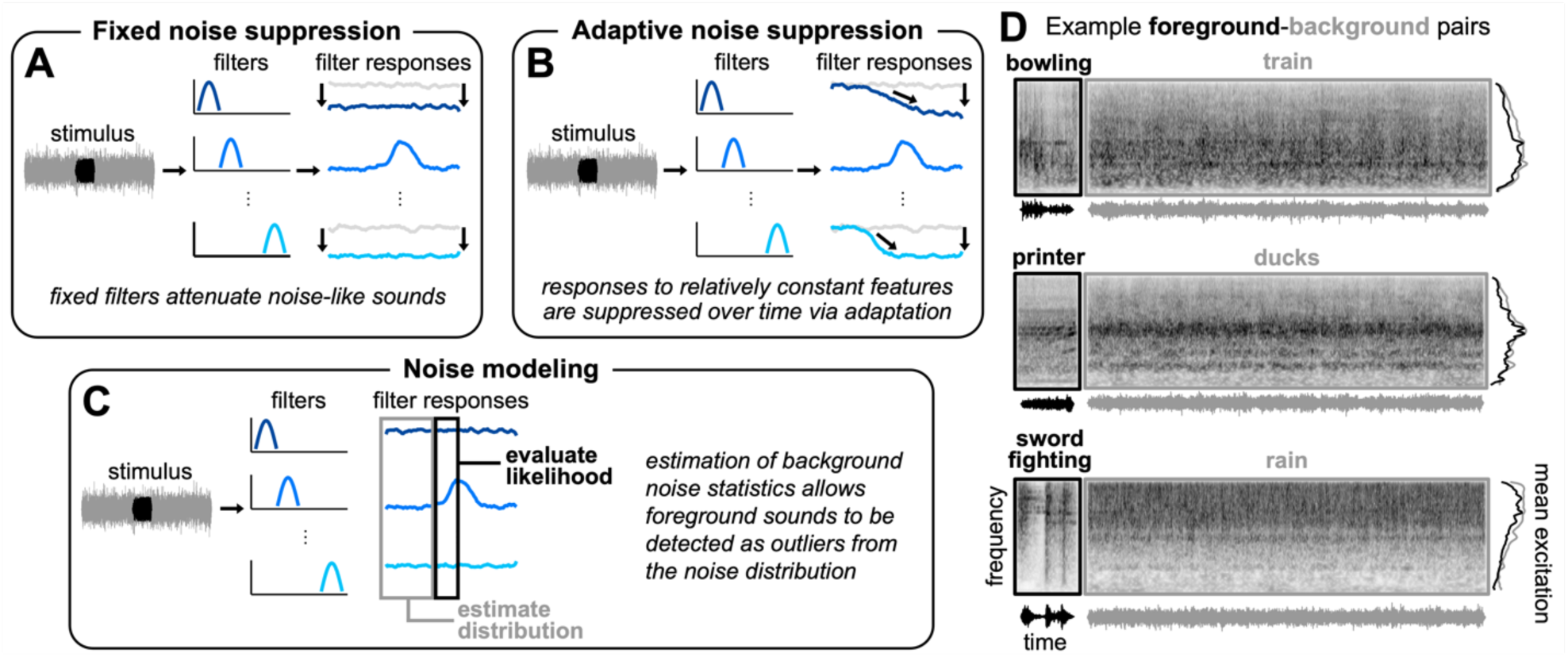
Potential explanations for noise-robust hearing and examples of experimental stimuli. (*A*) Fixed noise suppression hypothesis. A stimulus waveform (left, background shown in gray, foreground shown in black) passes through a set of filters (middle, transfer functions shown in shades of blue for an unspecified stimulus dimension), resulting in a set of filter responses over time (right). Here and in *B* and *C*, the filter tuning is unspecified and is not essential to the general predictions of the hypotheses. The filters could be nonlinear functions of the input and might measure higher-order properties of sound. In the fixed noise suppression hypothesis, the gain of a fixed set of filters is reduced to attenuate noise-like sounds (light gray responses show unattenuated filter response). (*B*) Adaptive noise suppression hypothesis. In the adaptive noise suppression hypothesis, responses to relatively constant features are suppressed over time via adaptation. (*C*) Noise modeling hypothesis. In the noise modeling hypothesis, estimation of background noise statistics allows foreground sounds to be detected as outliers from the associated distribution. (*D*) Example sounds used to generate experimental stimuli. Each panel shows the foreground (left, black) and background (right, gray) sound from an example trial, displayed as sound waveforms (bottom), cochleagrams (top), and mean excitation patterns (right). Cochleagrams were generated from the envelopes of a set of bandpass filters with tuning modeled on the human ear. Darker gray denotes higher intensity. Mean excitation patterns were obtained by averaging the cochleagram over time. In our initial experiments, foreground-background pairs were selected to have similar long-term spectra to minimize differences in the spectrotemporal overlap that would otherwise cause large variation in detectability from across pairs. This design choice turned out not to be essential and was dropped in later experiments.

An alternative possibility is that the auditory system might actively model the statistical structure of noise (Fig. 1C). This idea derives some a priori plausibility from the potential of background noise to convey useful information. Although laboratory studies of hearing in noise tend to use a single type of unstructured synthetic noise (e.g., white noise or pink noise), “noise” in the world can vary dramatically from place to place, often providing behaviorally relevant information about the environment (25, 26), such as the intensity of rain or wind. Such real-world noises are commonly referred to as auditory textures (24, 27). Auditory textures are typically generated by superpositions of many similar acoustic events and can exhibit a diversity of statistical properties (24). Moreover, human listeners are sensitive to statistical regularities of textures (24, 28–32) and estimate and represent their properties even in the presence of other sounds (33, 34). These considerations raise the possibility that rather than simply suppressing noise, the auditory system might model its statistical structure, using the resulting model to aid the separation of noise from other sound sources akin to how “schemas” are thought to aid the segregation of familiar words and melodies (35–37). Thus, there are at least three potential explanations for noise-robust hearing: fixed noise suppression, adaptive noise suppression, and the internal modeling of noise “schemas”.

We sought to test these three candidate explanations for noise-robust hearing and assess their role in everyday hearing. Adaptive suppression and internal noise modeling both require some form of online estimation of noise properties. Thus, both accounts predict that the ability to hear in noise should improve following the onset of a noise source, since its properties are more accurately estimated with larger samples. Such temporal effects have been documented in a few tasks (38) including pure tone detection (classically termed “overshoot”) (39, 40), amplitude modulation detection (41), phoneme recognition (18, 42) and word recognition (43, 44). However, because much of this work was conducted using relatively unstructured synthetic noise, it was unclear whether such temporal effects might be observed in more natural contexts (e.g., with realistic noise that is not fixed throughout a listening session). We thus began by characterizing listeners’ ability to detect, recognize and localize natural foreground sounds embedded in real-world background noise.

Although adaptive suppression and internal noise modeling are not necessarily mutually exclusive (see Discussion), they could be differentiated via the time course of their effects. Specifically, neural adaptation in the auditory system typically dissipates fairly rapidly following a stimulus offset (18), such that its effects would be expected to wash out during an interruption to background noise. By contrast, an internal model of noise properties might be maintained over time, yielding more persistent effects. Thus, to distinguish these two hypotheses, we further investigated whether noise robustness would persist across interruptions in noise, and whether it might improve following intermittent repeated exposure to particular background noises. Such improvement would be expected if listeners learn noise “schemas” akin to the schemas acquired for melodies (37), but not if they simply adapt to ongoing noise in the environment.

We found that the ability to detect, recognize, and localize foreground sounds in noise improved over the initial second of exposure to the background, a timescale substantially longer than previously reported for synthetic noise and artificial tasks. We also found that foreground detection performance was robust to temporary changes in the background, suggesting listeners maintain a representation of noise properties across interruptions. Moreover, detection performance was enhanced for recurring background noises, suggesting internal models of noise properties—noise schemas—are built up and maintained over time. Finally, we found that the pattern of human performance could be explained by an observer model that estimates the statistics of ongoing background noise and detects foreground sounds as outliers from this distribution. Taken together, the results suggest that the predictable statistical structure of real-world background noise is represented in an internal model that is used by the auditory system to aid hearing in noise.

## Results

To characterize real-world hearing-in-noise abilities, we sourced a diverse set of 160 natural foreground sounds and background noises to create experimental stimuli. Each stimulus consisted of a brief foreground sound paired with an extended background noise. Foreground sounds were 0.5 second excerpts of recorded natural sounds (45, 46), and background noises were 3.25 second excerpts of sound textures synthesized (24) from the statistics of real-world textures drawn from a large set of YouTube soundtracks (AudioSet) (47, 48). In our initial experiments, foreground-background pairs (Fig. 1D) were selected to have similar long-term spectra to avoid large differences in spectrotemporal overlap that would otherwise cause large variation in detectability from across pairs.

### Experiment 1: Foreground detection improves with exposure to background noise

We began by measuring the detection of natural sounds embedded in real-world background noises (Fig. 2A). On each trial, participants heard a continuous background noise presented either in isolation or with a brief “foreground” sound superimposed, then judged whether the stimulus contained one or two sound sources. Across trials, we manipulated both the temporal position and signal-to-noise ratio (SNR) of the foreground relative to the background. We chose the temporal positions so that there was always an onset asynchrony of at least 250 ms between foreground and background. Previously reported temporal dependencies of tone-in-noise detection (in which detection is better when tone and noise are asynchronous; “overshoot”) are limited to asynchronies of less than a few hundred milliseconds (39, 40, 49). However, it seemed plausible that longer timescales might be evident with more naturalistic noise sources and foreground sounds.

**Figure 2.**
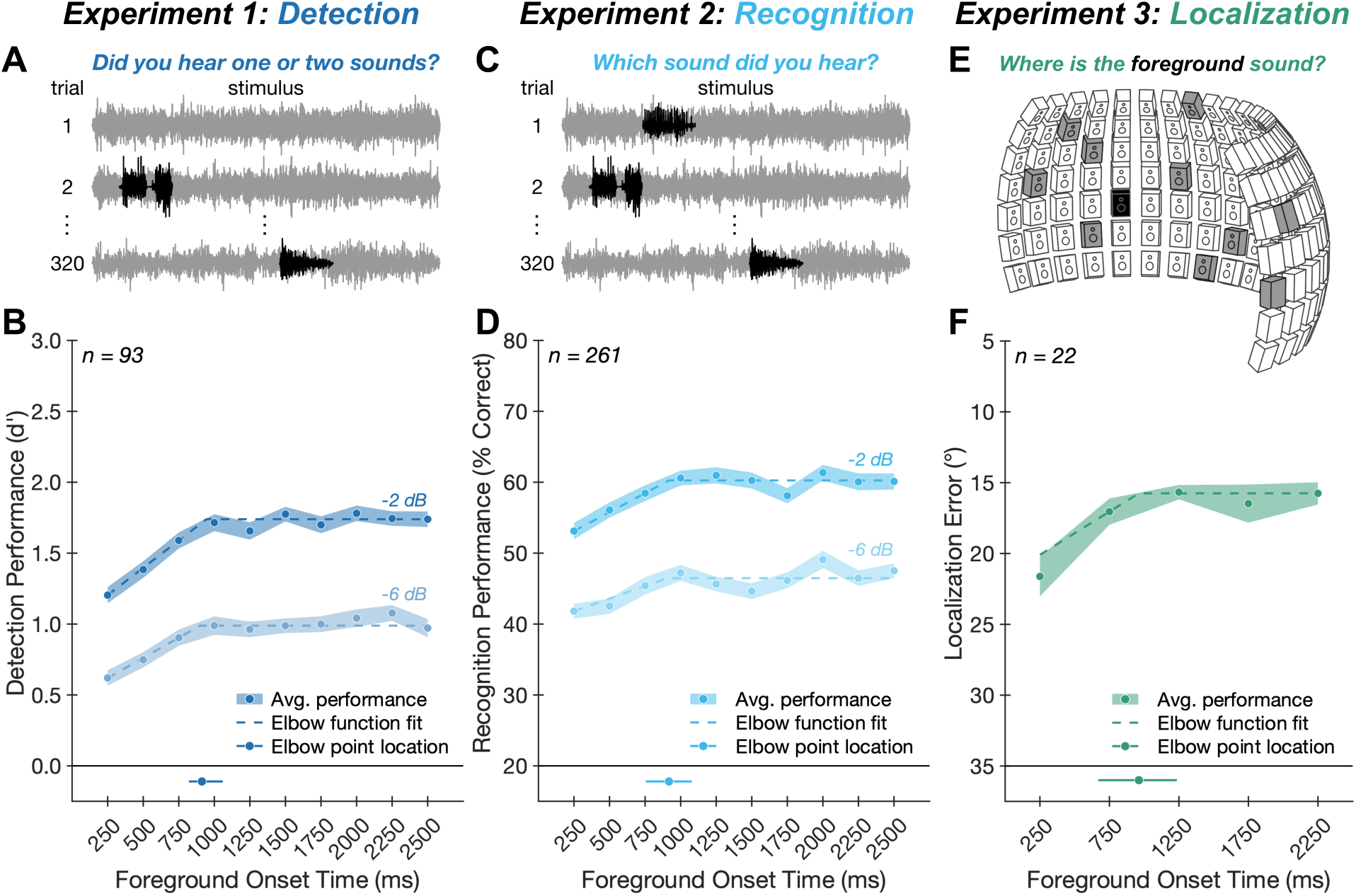
Experiments 1-3: Foreground detection, recognition, and localization improves with exposure to background noise. (*A*) Experiment 1 task. On each trial, participants heard a continuous background noise (gray) presented either in isolation (e.g., trial 1) or with a brief additional foreground sound (black) superimposed (e.g., trial 2). We manipulated the onset time and SNR of the foreground relative to the background. Participants judged whether the stimulus contained one or two sound sources. (*B*) Experiment 1 results. Average foreground detection performance (quantified as d’; blue circles) is plotted as a function of SNR and foreground onset time. Shaded regions plot standard errors. Dashed lines plot elbow function fit. Solid line below main axis plots one standard deviation above and below the median elbow point, obtained by fitting elbow functions to the results averaged over SNR and bootstrapping over participants; dot on this line plots the fitted elbow point from the complete participant sample. (*C*) Experiment 2 task. On each trial, participants heard background noise (gray) containing a foreground sound (black) and were asked to identify the foreground by selecting a text label from five options. (*D*) Experiment 2 results. Foreground recognition performance (quantified as percent correct; blue circles) is plotted as a function of SNR and foreground onset time. Chance performance was 20%. Data plotted using same conventions as *B*. (*E*) Experiment 3 task. Stimuli were presented via an array of 133 speakers spanning -90° to +90° in azimuth and -20° to +40° in elevation. On each trial, participants heard a scene composed of diffuse background noise (different samples of a texture played from 10 randomly selected speakers, shown in gray in the diagram) and a foreground sound (played from a randomly selected speaker, show in black in the diagram) occurring at one of five temporal positions within the noise. Participants judged the location of the foreground sound. (*F*) Experiment 3 results. Average foreground localization performance (quantified as absolute localization error in azimuth, in degrees; green circles) is plotted as a function of foreground onset time. The y-axis is oriented to match conventions in other panels where higher positions along the ordinate indicate better performance. Data plotted using same conventions as *B*.

Because task performance might benefit from knowledge of the foreground sounds (35–37), it was important that listeners only heard each foreground sound once during the experiment. To achieve this goal, we used an experimental design in which each participant completed only one trial for each of the 160 foreground-background pairings, with each pairing randomly assigned to one of the 20 experimental conditions (10 temporal positions crossed with 2 SNRs). Since this design necessitated a large sample size, we conducted this and other experiments online (with the exception of Experiment 3). Each of the 160 background noises also occurred once without a foreground sound. We calculated a single false-alarm rate from these background-only trials along with a hit rate for each of the 20 experimental conditions.

We found that foreground detection performance (quantified as d’) improved with exposure to the background (Fig. 2B; main effect of foreground onset time: F(9,828)=22.85, p<0.001, 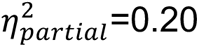). As expected, we also saw better foreground detection performance at the higher SNR (main effect of SNR: F(1,92)=769.15, p<0.001, 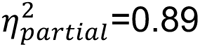), but the benefit of background exposure was evident at both SNRs (no significant interaction between foreground onset time and SNR: F(9,828)=0.76, p=0.65, 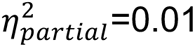). In both cases, task performance increased as the foreground sound was positioned later in the noise, with performance rising over roughly the initial second of exposure to the background.

This temporal dependence is consistent with the idea that listeners use the background noise preceding the foreground in order to perform the task. The temporal dependence also rules out several alternative possibilities. For example, if listeners performed the task entirely by detecting acoustic cues from the onset of the foreground sound, then task performance should be comparable at each temporal position of the foreground. Alternatively, if listeners could also perform the task equally well by listening retrospectively (using the background noise following the foreground to make a decision about the foreground’s presence), then performance should also be comparable across the different foreground positions, since the total duration of background noise is the same for each condition.

To better quantify the timescale of the effect, we fit an “elbow” function (a piecewise linear function consisting of two line segments; see Methods) to the results (averaged over SNRs). We bootstrapped over participants to obtain a confidence interval around the location of the elbow point (i.e., the transition from rise to plateau). This analysis indicated that foreground detection performance improved with exposure to the background before reaching a plateau after 912 ms (95% CI: [812, 1223] ms).

### Experiment 2: Exposure to background noise benefits sound recognition

In Experiment 2, we asked whether the benefit of background exposure extends to a recognition task. On each trial, participants heard a foreground-background pairing from Experiment 1 and were asked to identify the foreground by selecting a text label from five options (Fig. 2C). One option was the correct label; the remaining options were chosen randomly from the labels of the other foreground sounds in the stimulus set.

Recognition performance improved with exposure to the background in much the same way as did detection (Fig. 2D; main effect of foreground onset time: F(9,2340)=9.04, p<0.001, 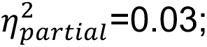 no significant interaction between foreground onset time and SNR: F(9,2340)=0.84, p=0.58, 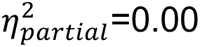). The elbow function fit to these results indicated a timescale of improvement similar to that in the detection task from Experiment 1, with a plateau in performance after 905 ms (95% CI: [726, 1242] ms) of exposure to the background.

### Experiment 3: Exposure to background noise benefits sound localization

We next asked whether the ability to localize sounds in noise similarly benefits from exposure to the background. We conducted this experiment in-lab using an array of speakers (Fig. 2E). On each trial, participants heard a scene composed of a foreground sound superimposed on spatially diffuse background noise, with the foreground occurring at one of five temporal positions within the background. Participants sat facing the array, holding their head still, and localized the foreground sound, entering the label of the corresponding speaker as their response. Because this experiment had to be run in-person (rather than online), we chose to use only five temporal positions at a single SNR in order to reduce the total number of conditions, thereby increasing power and allowing us to collect data from a modest number of participants. Additionally, we lowered the SNR to account for the likelihood that spatial cues would reduce detection thresholds (50). It turned out that at the tested SNR, localization in elevation was close to chance. Thus, we quantified sound localization performance using the absolute localization error in azimuth only.

Sound localization improved with exposure to the background in a manner similar to that observed for detection and recognition tasks (Fig. 2F; main effect of foreground onset time: F(4,84)=6.09, p<0.001, 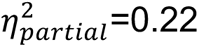), with performance plateauing after 962 ms (95% CI: [750, 1954] ms) of exposure to the background. Overall, the results point to a consistent benefit from noise exposure, spanning detection, recognition, and localization of natural sounds in noise.

### Experiment 4: Benefit of background exposure persists despite knowing what to listen for

In real-world conditions, we often listen for particular sounds in an auditory scene. For example, when crossing the street, one might listen out for crosswalk signals, bike bells, or accelerating engines. Because expectations about a source can aid its segregation from a scene (35–37), and might also benefit hearing in noise, it was unclear whether the benefit of background exposure would persist if participants knew what to listen for. To address this issue, we conducted a variant of Experiment 1 in which participants were cued to listen for a particular foreground sound on each trial (SI Appendix, Fig. S1A). On each trial, participants first heard a foreground sound in isolation (the “cue”), followed by continuous background noise. Half of the trials contained the cued foreground sound superimposed somewhere on the background, and participants judged whether the cued sound was present. We again found that foreground detection performance improved with exposure to the background (SI Appendix, Fig. S1B; main effect of foreground onset time: F(9,1215)=19.91, p<0.001, 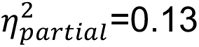). The timescale of improvement was similar to that in the detection task from Experiment 1, with a plateau in performance after 885 ms (95% CI: [582, 1766] ms) of exposure to the background (not significantly different from the elbow point in Experiment 1; p=0.52 via permutation test, elbow point difference: 68 ms). These results demonstrate that the benefit of background exposure persists even when participants know what to listen for, highlighting the relevance of this phenomenon for a range of real-world contexts.

### An observer model based on background noise estimation replicates human results

The results from Experiments 1-4 demonstrate a benefit of background noise exposure that provides evidence against the fixed noise suppression hypothesis, but that is conceptually consistent with both the adaptive suppression and internal noise modeling hypotheses. To first establish the plausibility of background noise estimation as an account of human hearing in noise, we built a signal-computable observer model to perform the foreground detection task from Experiment 1. The model evaluates the likelihood of incoming samples under a distribution whose parameters are estimated from past samples. The key idea is that samples not belonging to the background distribution (i.e., samples from the foreground sound) will tend to have low likelihood under a model of the background. Thus, the model can detect a foreground sound via samples that it assigns low likelihood. Intuitively, the fitted noise distribution should become more accurate with more samples, making it easier to detect outliers. However, it was not obvious that this type of model would achieve variation in performance with onset time that was on par with that observed in humans. We asked whether such a model could replicate the temporal dependence of foreground detection in noise observed in our human participants.

A schematic of the model is shown in Fig. 3A. First, an input sound waveform is passed through a standard model of auditory processing consisting of two stages: a peripheral stage modeled after the cochlea (yielding a “cochleagram”), followed by a set of spectrotemporal filters inspired by the auditory cortex that operate on the cochleagram, yielding time-varying activations of different spectrotemporal features. Next, a probability distribution is estimated from these activations over a past time window and used to evaluate the negative log-likelihood of samples in a “present” time window. This quantity (“surprisal”) measures how unexpected the present samples are given the learned background distribution. The process is then stepped forward in time and repeated, resulting in a set of surprisal values for each spectrotemporal filter at each time point within the stimulus. The surprisal is then averaged across filter channels and compared to a decision threshold to decide whether a foreground sound is present. The decision threshold was determined empirically as a value of surprisal substantially greater than would be expected by chance (see Methods).

**Figure 3.**
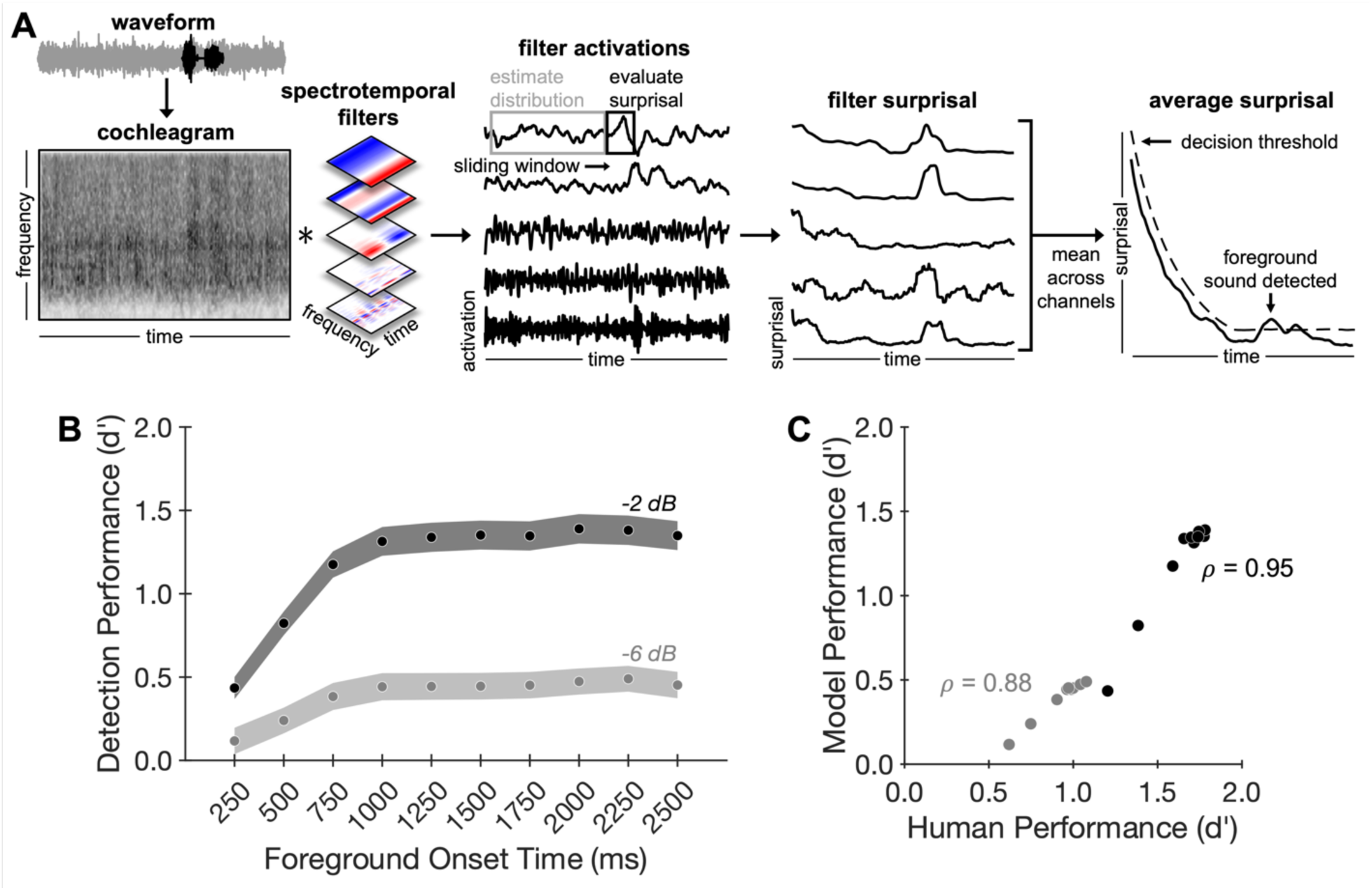
An observer model based on background noise estimation replicates human results. (*A*) Model schematic. First, an input sound waveform is passed through a standard model of auditory processing. This model consists of two stages: a peripheral stage modeled after the cochlea (yielding a cochleagram, first panel), followed by a set of spectrotemporal filters inspired by the auditory cortex that operate on the cochleagram, yielding time-varying activations of different spectrotemporal features (second panel). A sliding window is used to evaluate the negative log-likelihood (“surprisal”) within each filter channel over time (third panel). Finally, the resulting filter surprisal curves are averaged across channels and compared to a time-varying decision threshold to decide whether a foreground sound is present (fourth panel; y-axis is scaled differently in third and fourth panels to accommodate the surprisal plots for multiple individual filters). (*B*) Model results. Model foreground detection performance (quantified as d’) is plotted as a function of SNR and foreground onset time. Shaded regions plot standard deviations of performance obtained by bootstrapping over stimuli. (*C*) Human-model comparison. Model performance is highly correlated with human performance on the foreground detection task (Experiment 1) for both the -2 dB (black circles) and -6 dB (gray circles) SNR conditions.

For simplicity we implemented the model with univariate normal distributions fit to each filter output as these were sufficient to account for the qualitative effects seen in human judgments. We note that this choice results in an impoverished model of sound texture. In particular, the distribution only models the mean and variance of filter activations while ignoring other higher order statistics (e.g., correlations across filters) known to be important for sound texture perception (24). It also ignores temporal structure in the signal that exceeds the width of the filter kernels, treating all filter activations as independent. Although natural textures sometimes contain such high-order structure, it is at present unclear how this structure should be captured in a probabilistic model (existing models of texture (24) are based on a set of statistics rather than explicit probability distributions as are needed to evaluate outliers), and so we chose to sidestep this question to obtain a proof of concept for the general approach. The resulting simplifications would be expected to lower performance relative to what would be obtained with a distribution that more completely accounts for the statistical structure of natural textures. That said, the model captured some of the spectrotemporal structure of natural textures that differentiates them from traditional synthetic noise, and so seemed a reasonable choice with which to explore the general hypothesis of noise modeling.

The model was additionally defined by two hyperparameters: the width of the past window over which noise distribution parameters were estimated, and the width of the present window over which surprisal was averaged. We tested a range of past and present window sizes and found that the best match to human data occurred with a past window size of 1000 ms and a present window size of 500 ms. These results are presented here (see SI Appendix, Fig. S2 for the human-model correlation for different window sizes).

Despite its simplicity, the model qualitatively replicated the results from Experiment 1, showing a similar pattern of improvement with exposure to the background (Fig. 3B; see SI Appendix, Fig. S2 for model results with alternative window sizes, which remained qualitatively consistent with human results). Although model performance was below that of humans, the overall pattern of model performance across conditions was highly correlated with human performance (-2 dB SNR: *ρ*=0.95, p<0.001; -6 dB SNR: *ρ*=0.88, p=0.002). Overall, these results support background noise estimation as a plausible account of human hearing in noise by demonstrating that the qualitative trends evident in human behavioral performance can be explained by a model that estimates the statistics of ongoing background noise.

### Experiments 5a & 5b: Foreground detection is robust to background interruptions

Experiments 5a and 5b aimed to distinguish the adaptive suppression and noise modeling hypotheses by testing the effect of interruptions to the background. We modified the stimuli from Experiment 1, temporarily interrupting the background noise with either silence or white noise, initially using a 500 ms interruption (Experiment 5a; Fig. 4A). The rationale was that this change to the background might cause adaptation to “reset”, leading to a decrement in foreground detection performance following the interruption. By contrast, the noise modeling hypothesis could allow for a benefit from background exposure despite the interruption, because the estimated noise parameters could be stored across the interruption.

**Figure 4.**
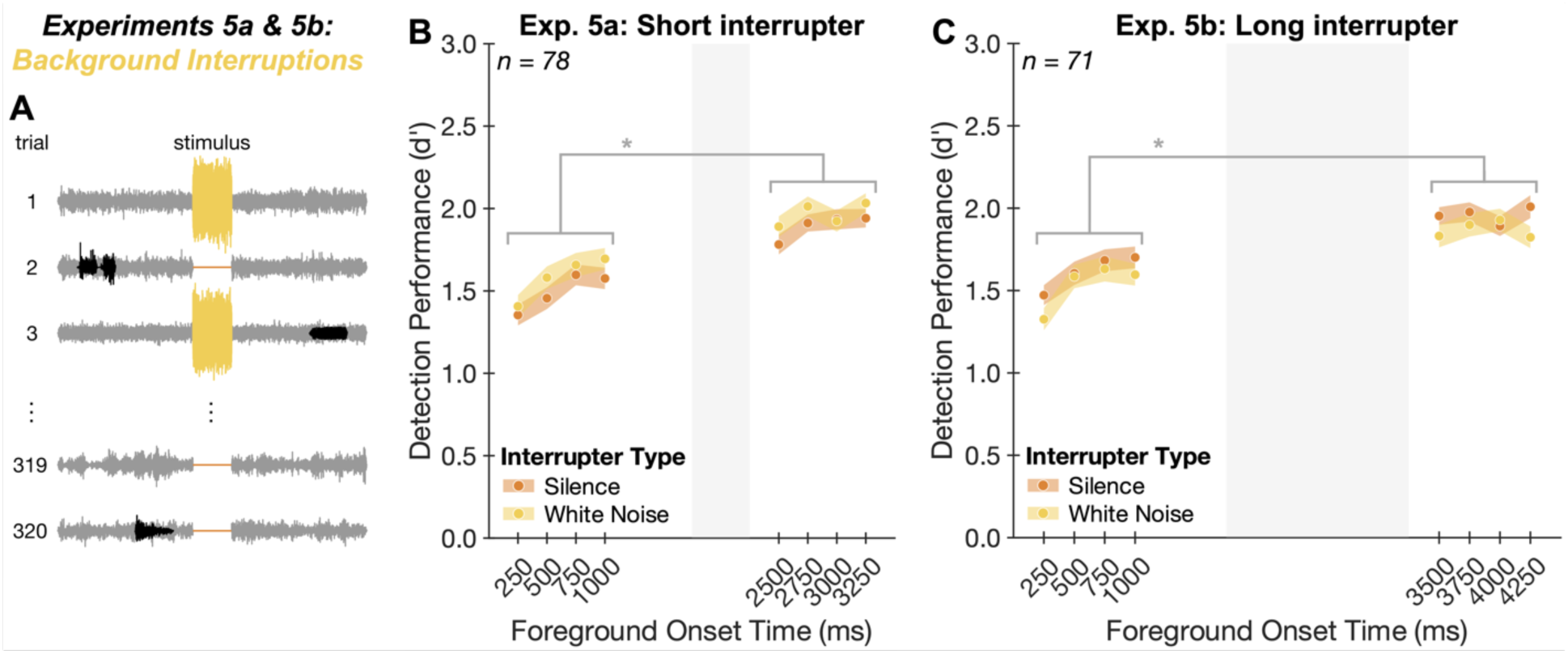
Experiments 5a & 5b: Foreground detection is robust to background interruptions. (*A*) Experimental task. Stimuli were like those from Experiment 1 but were modified by replacing the middle 500 ms (Experiment 5a) or 1500 ms (Experiment 5b) of background noise with either silence (orange) or white noise (yellow). Participants were asked to ignore this interruption and judge whether the stimulus contained one or two sound sources. (*B*) Experiment 5a results. Average foreground detection performance (quantified as d’) is plotted as a function of interrupter type and foreground onset time. Shaded regions plot standard errors. Gray region denotes temporal position of interruption in background noise. * indicates statistical significance, p<0.001. (*C*) Experiment 5b results. Same conventions as *B*.

Consistent with this latter possibility, foreground detection performance was greater for foregrounds following the interruption compared to those preceding the interruption (Fig. 4B; main effect of foreground position relative to interrupter: F(1,77)=172.86, p<0.001, 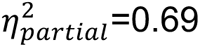). The pattern of results was similar for silent and noise interruptions (no significant interaction between interrupter type and foreground position relative to interrupter: F(1,77)=0.12, p=0.73, 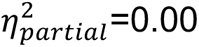).

To address the possibility that a 500 ms interruption was insufficient to trigger a complete release of adaptation (51), we ran an additional experiment (Experiment 5b) in which we increased the duration of the interrupter to 1500 ms and asked whether the benefit of background exposure persisted. Despite the longer interruption, we again found that detection performance was greater for foregrounds following the interruption compared to those preceding the interruption (Fig. 4C; main effect of foreground position relative to interrupter: F(1,70)= 134.48, p<0.001, 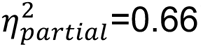). The pattern of results was again comparable for noise and silent interruptions (no significant interaction between interrupter type and foreground position: F(1,70)=0.01, p=0.92, 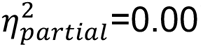).

Perhaps the clearest evidence against an adaptation explanation is the fact that the results appear to not be affected by the duration of the interruption (500 vs 1500 ms; no significant effect of interrupter duration when comparing performance for onset times after the interruption: F(1,147)=0.08, p=0.77, 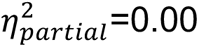). Although the parameters of any adaptive processes that might be at play are not definitively established, one would almost surely expect a difference in release from adaptation for differences in interruption durations of this magnitude. Taken together, the results of Experiments 5a and 5b indicate that the benefit of background exposure is unlikely to reflect adaptation alone. Instead, listeners appear to maintain an internal representation of noise properties across temporary interruptions.

### Experiments 6 & 7: Repetition of background noise enhances foreground detection

We next investigated whether internal models of noise are built up over time, akin to the “schemas” that can be learned for recurring patterns in speech and music (36, 37). If listeners learn noise schemas and use them to aid hearing in noise, foreground detection should be enhanced for frequently recurring background noises. It also seemed plausible that the learning of a schema might reduce the “delay benefit”—the improvement in performance as the foreground onset is delayed relative to the noise onset—since listeners could use a stored representation of the noise properties, rather than having to estimate them online. We thus also tested whether the delay benefit would be altered if the noise repeated across trials.

We first ran a variant of Experiment 1 in which a subset of the background noises (selected randomly for each participant) occurred repeatedly over the course of the experiment (Fig. 5A). A background noise was repeated on every trial in blocks of 40 trials, with each block containing a different repeating noise. We used unique noise exemplars for each repetition so that listeners would have to learn the statistical properties of the noise (28) rather than the specific exemplar (52, 53). We note that adaptation could, in principle, be expected to build up over the course of the block of repeated noise, potentially also accounting for altered performance. This experiment was thus not intended to distinguish noise schemas from adaptation, but rather to test a prediction of noise schemas in a simple setting before probing for benefits of repeated noises that might be less likely to be produced by adaptation (Experiment 8).

**Figure 5.**
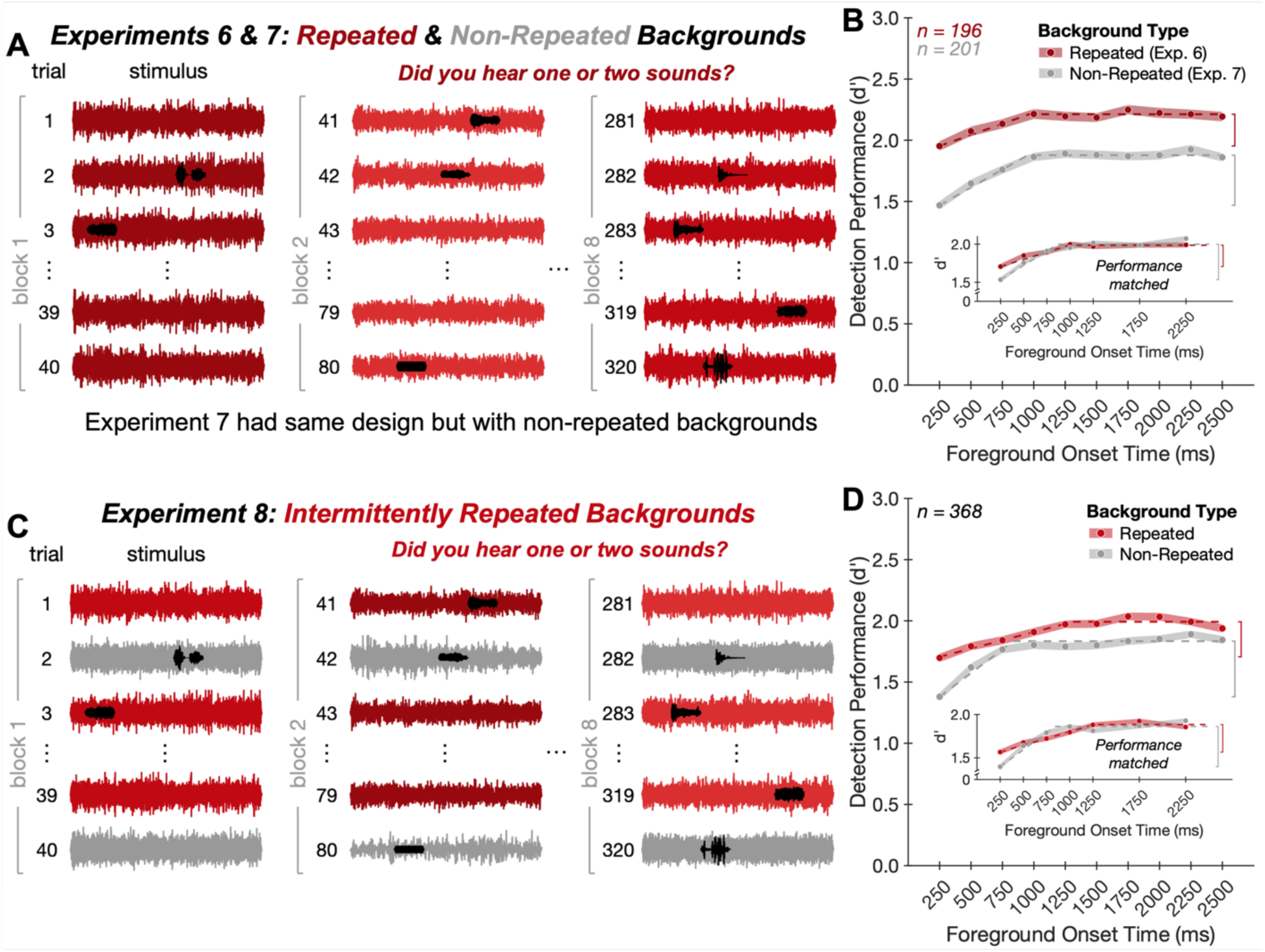
Experiments 6-8: Repetition of background noise enhances foreground detection. (*A*) Experiment 6 design. A background noise was repeated (red waveforms) on every trial in blocks of 40 trials, with each block containing a different repeating background noise (denoted by different shades of red). Participants judged whether the stimulus contained one or two sound sources. To ensure listeners would not benefit from learning the structure of the foregrounds, each foreground occurred only once with foregrounds and backgrounds paired randomly across participants. This design necessitated a companion experiment with similarly uncontrolled foreground-background pairings in which the backgrounds were not repeated across trials (Experiment 7; not shown). (*B*) Experiment 6 and 7 results. Average foreground detection performance (quantified as d’) is plotted as a function of foreground onset time for repeated (red circles) versus non-repeated (gray circles) backgrounds. Shaded regions plot standard errors. Dashed lines plot elbow function fit. Vertical brackets denote the delay benefit. Inset shows results after matching asymptotic performance across groups of participants. (*C*) Experiment 8 design. The experiment was identical to Experiment 6 except that background noises were repeated (red waveforms) on every other trial within a block with intervening trials containing non-repeating backgrounds (gray waveforms). (*D*) Experiment 8 results. Same conventions as *B*.

Each foreground sound occurred only once throughout the experiment to avoid the possibility that listeners might instead benefit from learning the structure of the foreground. As a result, we had to forego the controlled foreground-background pairings used in Experiments 1-5 and instead allowed foregrounds and backgrounds to be paired randomly across participants, while lowering the SNR to partially compensate for the decreased average spectral overlap between foreground and background. This design constraint necessitated a companion experiment (Experiment 7) with similarly uncontrolled foreground-background pairings in which the backgrounds varied across trials, with each of the 160 background noises occurring once with a foreground and once without a foreground, as in Experiment 1.

In the main analysis of interest, we found that performance was enhanced for repeating compared to non-repeating background noises (Fig. 5B; main effect of background type: F(1,395)=90.82, p<0.001, 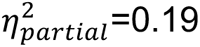). This enhancement developed over the course of a block in which the noises were repeated (SI Appendix, Fig. S3, main effect of first vs. second half of trials within a block: F(1,195)=17.44, p<0.001, 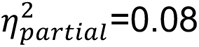). Although an effect of foreground onset time remained evident when the noise was repeated (main effect of foreground onset time for Experiment 6: F(9,1755)=10.75, p<0.001, 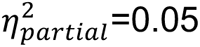), there was a significant interaction between foreground onset time and background repetition (F(9,3555)=2.52, p=0.01, 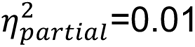). Specifically, the delay benefit was smaller for repeated backgrounds compared to non-repeated backgrounds (significant difference in the delay benefit; p<0.001 via permutation test, delay benefit difference: 0.15 in units of d’). To ensure the reduced delay benefit for repeating backgrounds was not driven by participants with near-ceiling performance, we ran a control analysis in which we selected groups of participants to have similar asymptotic performance using data from foreground onset times of 1500, 2000, and 2500 ms (see Methods), then measured the delay benefit using the data from the remaining foreground onset times for these participants. After matching asymptotic performance across groups of participants, the reduction in delay benefit persisted for repeated compared to non-repeated backgrounds (Fig. 5B inset, significant difference in delay benefit; p=0.01 via permutation test, delay benefit difference: 0.17 in units of d’).

Overall, these results confirm one prediction of the schema-based account of noise robustness: detection performance is improved for recurring backgrounds and less dependent on online noise estimation. We also note that these findings help reconcile the results in this paper with those of more traditional experimental paradigms, which repeat the same type of background noise throughout an experiment and find less pronounced temporal effects than those shown here.

We additionally note the results seem to be qualitatively unaffected by whether the foreground-background pairings were controlled. The effect of foreground onset time was similar in Experiment 7 (uncontrolled pairings) compared to Experiment 1 (controlled pairings), with no significant interaction between the experiment and the effect of foreground onset time (SI Appendix, Fig. S4; F(9,2628)=0.65, p=0.75, 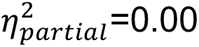). Both the timescale of improvement and the delay benefit were similar between the two experiments (no significant difference in elbow point: p=0.72 via permutation test, elbow point difference: 41 ms; no significant difference in delay benefit: p=0.50 via permutation test, delay benefit difference: 0.03 in units of d’).

### Experiment 8: Foreground detection is enhanced for intermittently repeated background noises

We next asked whether the benefit from recurring noises would be preserved across intervening stimuli, as might be expected if noise schemas are retained in memory, but not if the benefit reflects standard adaptation. In Experiment 8, the same type of background noise occurred on every other trial within a block (Experiment 8; Fig. 5C). As in Experiment 6, each block contained a different repeating background noise with unique noise exemplars for each repetition and each foreground sound was presented once with foregrounds and backgrounds paired randomly across participants.

We again found that performance was enhanced for repeating compared to non-repeating background noises (Fig. 5D; main effect of background type: F(1,367)=63.03, p<0.001, 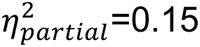). Additionally, there was again a significant interaction between the effect of foreground onset time and whether the background was repeated or not (F(9,3303)=4.38, p<0.001, 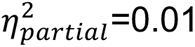), such that the delay benefit was smaller for repeating compared to non-repeating backgrounds (significant difference in delay benefit: p=0.002 via permutation test, delay benefit difference: 0.17 in units of d’). This difference persisted after matching asymptotic performance across subsets of participants (Fig. 5D inset, significant difference in delay benefit; p=0.03 via permutation test, delay benefit difference: 0.13 in units of d’). These results suggest that noise schemas— representations of noise statistics that aid foreground detection—are built up over time, maintained across intervening stimuli, and lessen the dependence on online noise estimation.

### Experiment 9: Benefit of background exposure is reduced for stationary noise

Across multiple tasks, we consistently found an improvement in performance over the initial second of exposure to the background. What factors might govern the timescale of this effect? A model that estimates statistics within an integration window (as in the model of Fig. 3) should exhibit improved performance up to the window width. One intuitive possibility is that the window width reflects a tradeoff between obtaining a good estimate of the background noise statistics (better for longer windows) and being able to resolve changes in these statistics (better for shorter windows). However, the accuracy with which statistics can be estimated for a given window size depends on the stability of the noise statistics over time (i.e., the stationarity of the noise). This observation raises the possibility that the optimal estimation window could be shorter for more stationary noise. To test whether these considerations might influence hearing in noise, we modified the stimuli from Experiment 1, replacing the real-world texture backgrounds with spectrally matched noise (Fig. 6A) to create noise backgrounds with increased stationarity. We quantified stationarity with a measure of the standard deviation of texture statistics across time windows (17, 33, 34, 48) (see Methods) and confirmed the increase in stationarity for spectrally matched noise backgrounds (Fig. 6B). Because the detection task is easier with more stationary noise, we reduced the SNRs of the foreground relative to the background to avoid ceiling performance.

**Figure 6.**
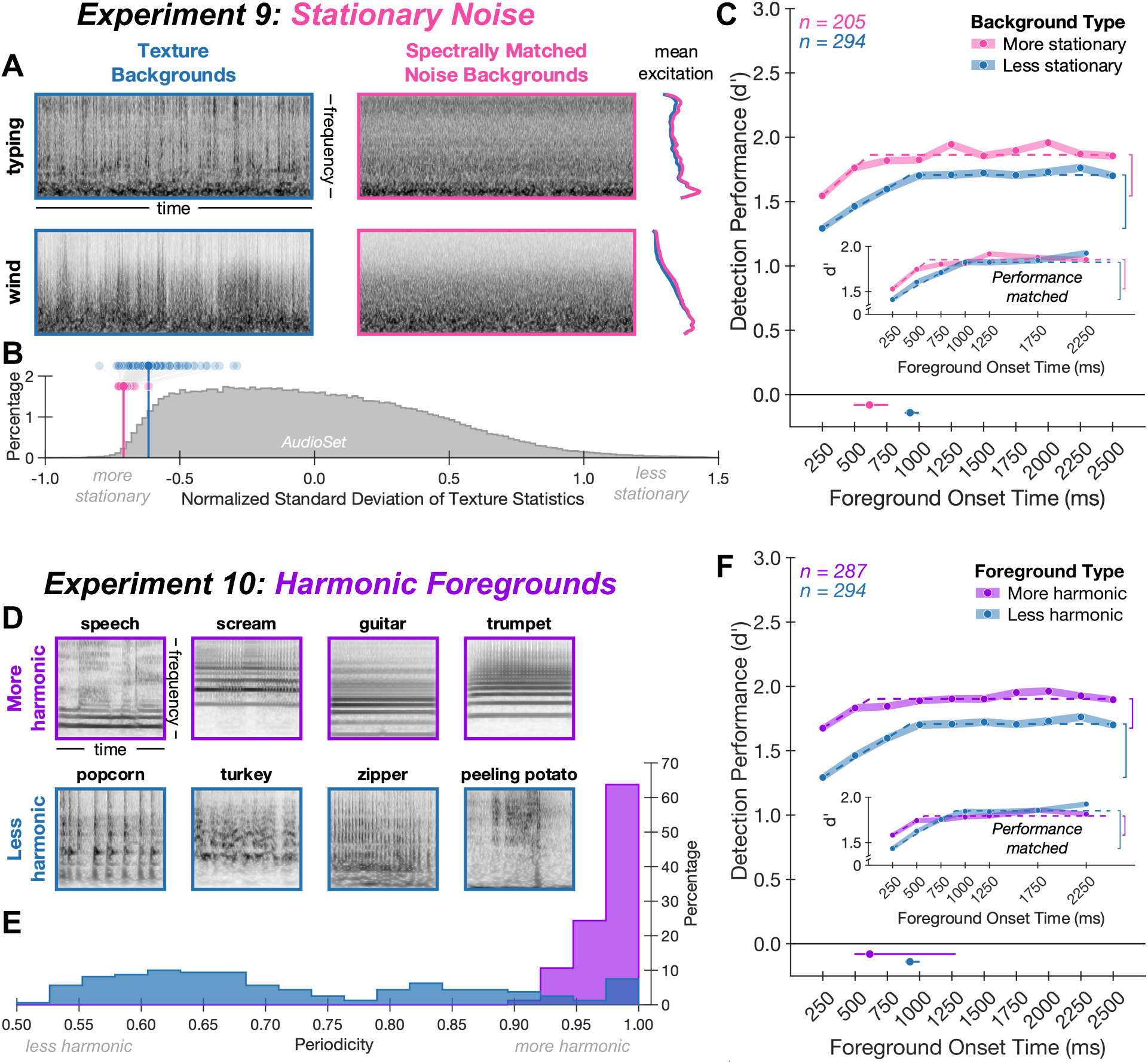
Experiments 9 & 10: Benefit of background exposure is reduced for stationary noise and harmonic foregrounds. (*A*) Example background noises from Experiment 9. The real-world texture backgrounds (left, blue) used in Experiments 1-8 were replaced with spectrally matched noise (middle, pink) to increase the noise stationarity. Backgrounds are displayed as cochleagrams (with darker gray indicating higher intensity) and mean excitation patterns (right). (*B*) Stationarity of background noises. Shaded circles indicate a measure of stationarity (standard deviation of texture statistics over time, normalized to account for increased variability of some statistics relative to others; see Methods) for the texture backgrounds used in Experiments 1-8 (shown in blue) and the spectrally matched noise backgrounds used in Experiment 9 (shown in pink). Gray lines connect textures to their spectrally matched counterparts, illustrating that the spectrally matched noise is generally more stationary than its texture counterpart. Vertical lines indicate mean stationarity of background noises in each stimulus set. For comparison, a histogram of stationarity scores calculated from a large set of YouTube soundtracks (AudioSet; see Methods) is shown in dark gray. Both sets of background noises are more stationary than the average soundtrack. (*C*) Experiment 9 results.

As in our previous experiments, foreground detection performance improved with exposure to the background (Fig. 6C; main effect of foreground onset time: F(9,1836)=25.99, p<0.001, 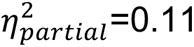). However, the effect was more modest than that observed with more naturalistic noise (significant interaction between foreground onset time and stationarity: F(9,3636)=2.69, p=0.004, 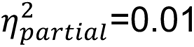). In particular, the timescale of improvement was shorter for stationary noise backgrounds compared to texture backgrounds (significant difference in elbow point for Experiment 9 compared to pooled results from Experiments 1 and 7: p=0.01 via permutation test, elbow point difference: 319 ms), and the delay benefit was reduced (significant difference in delay benefit: p=0.01 via permutation test, delay benefit difference: 0.10 in units of d’). This reduction in delay benefit remained after matching asymptotic performance across groups of participants (Fig. 6C inset, significant difference in delay benefit; p=0.03 via permutation test, delay benefit difference: 0.11 in units of d’). These findings help to further reconcile the results of this paper with prior work that has predominantly used highly stationary synthetic noise and has found weaker effects of onset time. The temporal effects we nonetheless observed could reflect the fact that the background noise spectrum varied from trial to trial in our experiments (unlike most experiments in prior work).

Average foreground detection performance (quantified as d’) is plotted as a function of foreground onset time for more stationary (pink circles) versus less stationary (blue circles; obtained from pooled results of Experiments 1 and 7) backgrounds. Shaded regions plot standard errors. Dashed lines plot elbow function fit. Vertical brackets denote the delay benefit. Solid lines below main axis plot one standard deviation above and below the median elbow points, obtained by bootstrapping over participants; dots on these lines plot the fitted elbow points from the complete participant samples. Inset shows results after matching asymptotic performance across groups of participants. (*D*) Example foreground sounds from Experiment 10. The foreground sounds used in Experiments 1-9 (bottom, blue) were replaced with human vocalizations and musical instrument sounds (top, purple). Foregrounds are displayed as cochleagrams. (*E*) Harmonicity of foregrounds. Harmonicity was quantified with a measure of waveform periodicity (see Methods). Histograms of periodicity are shown for the set of human vocalizations and instrument sounds used in Experiment 10 (purple) and for the set of foregrounds used in Experiments 1-9 (blue). (*F*) Experiment 10 results. Average foreground detection performance (quantified as d’) is plotted as a function of foreground onset time for more harmonic (purple circles) versus less harmonic (blue circles; obtained from pooled results of Experiments 1 and 7) foregrounds. Conventions same as *C*.

As with other effects of stationarity on integration timescales (33), there are at least two computational accounts of these results. One is that there is a single statistical estimation window that changes in temporal extent depending on the input stationarity. Another is that there are multiple estimation windows operating concurrently (potentially estimating different statistical properties), with a decision stage that selects a window (or combination of windows) on which to base responses. For instance, by selecting the shortest window that enables a confident decision, a decision stage might determine that a shorter estimation window is most appropriate for stationary noise.

### Effect of background exposure depends on foreground-background similarity

The possibility of concurrent estimation windows of different extents raised the question of whether the similarity of the foreground to the background could also influence the timescale of the effect of background exposure. Specifically, foregrounds that differ more noticeably from a background might be detectable with a less accurate model of the background that could be estimated with fewer samples (via a shorter estimation window). As one test of this possibility, we re-analyzed the results of Experiment 7, dividing trials into two groups whose foreground-background pairings differed in spectrotemporal similarity. We found a significant interaction between foreground onset time and spectrotemporal similarity (SI Appendix, Fig. S5; F(9,1791)=3.90, p<0.001, 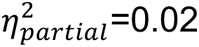), whereby the timescale of the noise exposure benefit was shorter for foreground-background pairs with lower spectrotemporal similarity (significant difference in elbow point: p=0.003 via permutation test, elbow point difference: 526 ms). This result is consistent with the idea that there are multiple concurrent windows for estimating noise statistics, with shorter windows being used when they are sufficient for a decision. The result also provides further evidence that the estimation of noise properties aids the detection of the foreground. In particular, the result helps rule out the possibility that the effect of onset time reflects interference between the processing of the background and the detection of the foreground (as could in principle happen if the noise onset initiated some involuntary process that was not used to detect the foreground and that instead initially impaired processing of concurrent sounds).

### Experiment 10: Benefit of background exposure is reduced for harmonic foregrounds

Motivated by the effect of foreground-background similarity, in the final experiment we tested the effect of background exposure on the detection of approximately harmonic foregrounds. We avoided human vocalizations and music instrument sounds in our initial experiments on the grounds that they are more detectable in noise (5) compared to other sounds and so could have introduced another source of variation in detection performance. However, the analysis of foreground-background similarity suggested that harmonic sounds might produce weaker effects of background exposure; given their prevalence in both everyday life and prior auditory experiments, this seemed important to test. The experiment was identical to Experiment 7 except that the foreground sounds used in previous experiments were replaced with excerpts from human vocalizations and musical instrument sounds (Fig. 6D). We confirmed that these sounds were more harmonic than those used in the previous experiments, using a measure of waveform periodicity (54) (Fig. 6E; see Methods).

A benefit of background exposure was evident for these (approximately) harmonic sounds (Fig. 6F; main effect of foreground onset time: F(9,2574)=13.66, p<0.001, 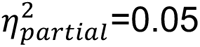), but it was weaker than that observed for less harmonic sounds (significant interaction between foreground onset time and harmonicity: F(9,4374)=4.25, p<0.001, 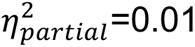). Compared to the pooled results from Experiments 1 & 7, the elbow point was earlier (significant difference in elbow point: p=0.03 via permutation test, elbow point difference: 317 ms) and the delay benefit was smaller (significant difference in delay benefit: p<0.001 via permutation test, delay benefit difference: 0.19 in units of d’). This difference persisted after matching performance across experiments (Fig. 6F inset; significant difference in delay benefit: p<0.001 via permutation test, delay benefit difference: 0.20 in units of d’). The results again help to reconcile our findings with previous work using speech or tones that have found smaller effects, while also showing that the qualitative effects of background exposure remain evident with harmonic foregrounds.

## Discussion

We investigated whether internal models of environmental noise are used by the auditory system to aid the perception of natural foreground sounds in background noise. We found that the ability to detect, recognize, and localize foreground sounds in noise improved over the initial second of exposure to the background. This benefit of background exposure persisted even when participants knew the foreground sound they had to listen for. The benefit of prior noise exposure was robust to temporary changes in the background and was enhanced for recurring backgrounds, suggesting that noise “schemas” are built up and maintained over time. We found that an observer model designed to capture the statistics of ongoing background noise could account for the pattern of human behavioral performance observed in the foreground detection task. Finally, we found evidence for a window of noise estimation that varies depending on the stimulus characteristics, appearing shorter both for more stationary noise and when foreground sounds are sufficiently distinct (e.g., by virtue of being harmonic) from the background so as to not require a detailed model of the background properties. Overall, the results suggest that the auditory system leverages internal models of noise properties—noise schemas—to facilitate the estimation of other concurrent sounds and support noise robust hearing.

### Relation to prior work

We sought to distinguish between three candidate explanations for noise robust hearing: fixed noise suppression, adaptive noise suppression, and internal modeling of noise “schemas”. Two lines of evidence have previously been presented in support of adaptive suppression (and contra fixed suppression). The first involves behavioral improvements in hearing abilities following exposure to a noise source (see (38) for a review), with improvements occurring over approximately 500 ms. A second body of research involves evidence of neural adaptation to noise (13, 14, 16, 18, 23), with related modeling work suggesting these adaptive responses could be explained by a mechanism that dynamically suppresses noise (23).

We built on this prior work in three respects. First, we found that foreground detection performance was robust to interruptions in the background and was enhanced for frequently recurring backgrounds. These findings are inconsistent with conventional adaptation to ongoing noise and instead suggest noise properties are estimated and maintained over time. Second, we demonstrated that the benefits of noise exposure on behavior generalize to natural stimuli and everyday listening contexts. In these conditions, behavioral performance improved over a period roughly twice as long previously reported, with performance plateauing around one second. These large effects appear to partly reflect the use of stimuli that vary across trials, realistic sources of noise, and diverse foreground sounds. We found smaller effects when noises repeated within an experiment (Experiments 6 & 8), when noise was more stationary (Experiment 9), and when foregrounds were harmonic (Experiment 10). These results reconcile our findings with previous work, which has tended to use a single type of highly stationary synthetic noise and harmonic foreground sounds, and which has seen smaller effects of time. Our results highlight the utility of assessing perception using natural stimuli, as it can reveal effects not fully evident with simpler traditional stimuli (55–57). Third, our experiments show that the temporal effects of exposure to background noise occur across multiple auditory tasks: detection, recognition, and localization.

The temporal dynamics of human task performance in noise could be explained by an observer model that estimates the statistics of ongoing background noise and detects foreground sounds as outliers from this distribution. This finding demonstrates that the observed improvement in task performance following noise exposure can result directly from a model of noise properties, and raises the question of how to reconcile our results with neurophysiological findings of noise suppression in the auditory system. We suggest that noise modeling and noise suppression are not mutually exclusive. One possibility is that the auditory system maintains parallel representations of sound: one in which noise properties are estimated and maintained, and another in which noise is suppressed to yield a relatively invariant representation of the foreground. The first representation could be used to derive the second, such that as noise becomes more accurately estimated (e.g., with more exposure to the background), the foreground representation becomes enhanced, as we found here. The existence of such parallel representations is consistent with neurophysiological findings that noise stimuli are represented subcortically (58, 59) and preferentially drive neurons in primary auditory cortex (60–62) but appear to be suppressed in non-primary auditory cortex (17, 63).

### Limitations

The model presented here provides evidence that estimation of noise statistics could underlie aspects of hearing in noise, but in its present form is not a complete account of human perception in this setting. As noted earlier, we modeled noise with relatively simple distributions that do not completely capture the structure known to be present in real-world noise. Although the approach was sufficient for our purposes, more sophisticated models will be required to fully account for human performance. A complete model would also estimate the properties of any foreground sounds in addition to detecting their presence. Intuitively, one might adopt an “old plus new” (64) approach in which samples that deviate from the distribution of ongoing background noise are interpreted as a (“new”) foreground sound whose features can be estimated as the “residual” after accounting for the background noise properties. The model as implemented here also does not account for the enhanced foreground detection observed for interrupted or frequently recurring backgrounds (Experiments 5-8). However, some of these effects could potentially be modeled by incorporating a prior over noise properties that is continually updated over the course of the experiment.

### The role of texture in auditory scene analysis

Using real-world noise signals, we found that the ability to hear in noise improves over the initial second of exposure to the background noise—substantially longer than the timescale previously reported for analogous tasks with simpler experimental stimuli. This relatively long timescale is broadly consistent with the growing literature on sound texture perception. Sound textures are thought to be perceptually represented in the form of time-averaged summary statistics (28) computed using averaging mechanisms with a temporal extent that depends on the texture stationarity (33) but is generally on the order of seconds. Moreover, the detection of changes in texture statistics documented in previous studies improves across a temporal scale similar to that observed in our work (29, 30). Other experiments have indicated that texture properties are estimated separately from other sound sources (33) and are filled in when masked by other sounds (34). Our results provide further evidence that sound texture plays a critical role in auditory scene analysis, as its estimation benefits the detection, recognition, and localization of other concurrent sounds. This growing literature supports the idea that background noise properties are actively estimated by the auditory system even in the presence of other sound sources.

### Reconsidering the role of noise

Sound textures are ubiquitous in everyday listening, constituting the background noise of many real-world auditory scenes. Yet research on hearing in noise has devoted relatively little attention to the role of noise itself. We consider hearing in noise as a form of auditory scene analysis in which listeners must segregate concurrent foreground and background sources from one another. From this perspective, noise is another source to be estimated rather than suppressed (65). However, the segregation of multiple sound sources is only possible because of the statistical regularities of natural sounds. Previous work has shown that human listeners can quickly detect repeating patterns in the acoustic input and use this structure to facilitate source segregation in artificial auditory scenes (66–68). The present results complement these findings by showing that the predictable statistical structure of noise is used to aid source segregation in natural auditory scenes.

### Future directions

Reverberation is another element of sound often thought of as a distortion that the auditory system must suppress to improve hearing in acoustic environments characteristically encountered in daily life (15, 69–71). It is analogously possible that the estimation, rather than suppression, of reverberation might help to recover the underlying sound source (72, 73). Thus, robustness to reverberation may be aided by an internal model of the statistical regularities that characterize real-world reverberation (72). “Schemas” of room reverberation may also be built up through short-term exposure to room-specific reverberation (74–77).

The computational principles described here are equally relevant to other sensory modalities. For instance, the detection or recognition of an object amid a cluttered visual scene can be viewed as a visual analogue of our hearing-in-noise experiments. As in hearing, the ability to visually recognize objects is impaired by clutter—a widely studied phenomenon known as visual crowding (78). Given that visual textures are thought to be represented by summary statistics averaged over space (79, 80), visual object recognition might be expected to improve with the size of a background texture region as background properties should be better estimated across larger spatial extents. Some preliminary evidence supports this hypothesis. Several studies from the visual crowding literature demonstrate a release from crowding when additional distractors or flankers are added to a display (81, 82). However, these studies are limited to relatively simple displays, and it remains to be seen whether such effects may be observed in more naturalistic settings.

## Materials and Methods

The code and data are available at https://github.com/mcdermottLab/NoiseSchemas.

### Stimulus selection

#### Foreground sounds

To select foreground sounds for our experimental stimuli, we began with a set of 447 2-second-long recordings of natural sounds used in previous experiments from our laboratory (45, 46). Because harmonicity can aid hearing in noise (5), we manually screened these recordings to remove any sounds containing music, speech, or human vocalizations (e.g., screams or grunts). Some approximately harmonic sounds nonetheless remained in the stimulus set (e.g., alarms, various beeping electronics, animal vocalizations, etc.); see Fig. 6E. Additionally, we removed texture-like sounds (e.g., hairdryer or crumpling paper) to help ensure that selected foregrounds would be distinct from the sound texture backgrounds they were superimposed on. This left us with a set of 167 foreground sounds. These foreground sounds were used for Experiments 1-9. Experiment 10 used a different set of foreground sounds chosen to be approximately harmonic (see “Experiment 10” below).

#### Background sounds

To select background sounds for our experimental stimuli, we screened a large set of audio examples (AudioSet (47)) for sound textures. Specifically, we first screened the “unbalanced train set” within AudioSet by excluding 1) any sound whose label indicated the presence of speech or music (e.g., “whispering”, “song”, etc.; see Table S1 for list of excluded labels), 2) any sound from the “sourceless” branch of the ontology, 3) any sound less than 10 s in length, and 4) any sound with greater than 1% of values equal to zero. This resulted in a set of 222,560 sounds (which were used for computing the normalization values used in the stationarity measure described below). We then computed a measure of stationarity developed in previous work from our lab (17, 33, 34) for each sound within this set and excluded any sound with a stationarity score above 0, leaving us with a large set of 142,922 AudioSet “textures.” From these AudioSet “textures”, we selected relatively stationary sounds by keeping sounds with stationarity scores between -0.75 and -0.67 (approximately the 87^th^ and 94^th^ percentiles of the sounds with scores below 0). Additionally, we sought to avoid periodic textures (e.g., rhythmic clapping or waves crashing) because foreground detectability within such textures greatly depends on the timing of the foreground relative to the period of the background texture. We measured the periodicity of each AudioSet texture as in previous work (34) by measuring the normalized auto-correlation of the envelope of the stimulus waveform and selecting the maximum peak between 125 ms and 500 ms (2–8 Hz). We kept only those sounds whose periodicity fell within 0.05 and 0.075 (approximately the 1^st^ and 7^th^ percentiles across all AudioSet textures). The intersection of the stationary and non-periodic textures yielded 1511 textures. We note that the stationarity analyses that were subsequently performed in this paper used a slightly different stationarity measure than the one used for the initial screening described above (the new measure was similar in spirit to the old one, but used a different form of normalization). As a result, the stationarity scores referenced above differ slightly from those reported in Fig. 6B.

The background noises used in our experiments were textures synthesized from statistics measured from the (recorded) AudioSet sound textures. There were two reasons for this choice. First, textures recorded in natural environments often contain distinctive acoustic events arising from other sources. For example, a recording of a stream might contain faintly audible bird calls. Such additional sources would create confusion in the experiments involving detection or recognition. Second, in several experiments (Experiments 3, 6 and 8) we needed to present multiple exemplars of the same texture, and for this purpose required more than 10 s of audio. For each of the 1511 textures, we created 9-second-long synthetic exemplars using a standard texture synthesis method (24). We found that the synthesis procedure converged (average SNR of all statistic classes was 20 dB or higher) for 1285 textures and selected the background noises from this set. We drew 3.25 s excerpts from these 9-second-long synthesized textures (see Foreground-background pairings below) to use in Experiments 1, 2, 4, 5, 7 and 10.

#### Stationarity measure

To quantify the stationarity of a sound for the analysis of Experiment 9, we computed a measure based on the standard deviation of texture statistics (24) across successive time windows (17, 33, 34, 48), based on the idea that stationary sounds have temporally stable statistical properties. Specifically, we first computed a set of texture statistics (subband mean, envelope mean, envelope standard deviation, envelope skew, envelope correlations, modulation band power, C1 modulation correlations, and C2 modulation correlations) for successive segments of a signal (using excerpt lengths of 0.125, 0.25, 0.5, 1, and 2 s). To put each of the statistics on the same scale, we then z-scored each statistic using the mean and standard deviation of each statistic calculated across the set of screened full-length AudioSet sounds (see “Background sounds” section above). To quantify how much each statistic changed across excerpts, we computed the standard deviation of each z-scored statistic across all excerpts of the same length. Because some statistics are intrinsically more variable than others, we computed a normalized measure of the variability of each statistic by dividing the computed standard deviations (separately for each statistic and excerpt length) by the average (i.e., expected) standard deviation across all the screened AudioSet sounds. To obtain a single measure of stationarity, we then averaged these normalized standard deviations across statistics and excerpt lengths. However, because some statistics classes contain more statistics than others, we first averaged the normalized standard deviations across all statistics within each class before averaging across the statistics classes (effectively weighting each statistics class equally) and excerpt lengths. The result is a normalized measure of statistic variability where smaller (i.e., more negative) values indicate greater stationarity.

#### Foreground-background pairings

In our initial experiments, experimental stimuli were generated from pairs of foregrounds and backgrounds selected to have similar long-term spectra to avoid large differences in foreground detectability across different foreground-background pairings. To select these pairs, we first created cochleagrams for each possible foreground and background sound. Cochleagrams were generated from the envelopes of a set of 38 bandpass filters (plus one low-pass and one high-pass channel) at a sampling rate of 500 Hz with tuning modeled on the human ear (24). Next, for each 2-second-long foreground sound, we randomly selected 100 0.5s cochleagram segments (from the entire 2s sound) and computed the Mahalanobis distance (*D*) between each foreground cochleagram segment and every background cochleagram. Specifically, we calculated the Mahalanobis distance for each point in time of the foreground cochleagram using the background cochleagram as the reference distribution, then averaged these distances over time: 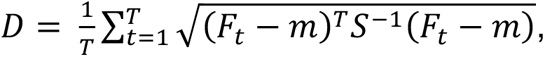 where *F_t_* is one column of the foreground cochleagram at time *t*, *m* is the excitation pattern (time-averaged cochleagram) of the background, and *S* is the covariance of the background cochleagram. The Mahalanobis distance quantifies the difference between the foreground and background excitation patterns while accounting for the covariance structure among cochlear channels measured from the background. For every possible foreground-background pair, we stored the foreground segment with minimum Mahalanobis distance then used the Hungarian algorithm (83)) to pair each of the 167 foregrounds with a background sound such that the Mahalanobis distance across the pairings was minimized. Finally, we manually listened to each of the selected background textures and selected 3.25 s excerpts that subjectively sounded fairly uniform. We then selected 7 of these pairings to use as practice trials, leaving the remaining 160 foreground-background pairings to be used as experimental stimuli. Table S2 lists each of the foreground-background pairings; the sound waveforms for each foreground and background are provided in the data and code repository for this paper.

### Experimental procedure for online participants

The condition-rich design of our experiments (e.g., 20 experimental conditions in Experiment 1), resulted in obtaining relatively few trials per condition per subject. To obtain the large sample sizes necessary to attain reliable results, we conducted our experiments (with the exception of Experiment 3) online using the Amazon Mechanical Turk and Prolific crowdsourcing platforms. Experiments 1 and 4 were conducted in 2021-2022 on Amazon Mechanical Turk. Experiments 2 and 5-10 were conducted in 2023-2024 on Prolific. Across multiple studies from our laboratory, we have found that online data can be of comparable quality to data collected in the lab provided a few modest steps are taken to standardize sound presentation, encourage compliance and promote task engagement (9, 34, 37, 57, 84, 85).

All participants provided informed consent and the Massachusetts Institute of Technology Committee on the Use of Humans as Experimental Subjects (COUHES) approved all experiments. Amazon Mechanical Turk participants were required to be in the United States, to have a HIT approval rate of greater than 95%, and to have had more than 100 HITs approved. Prolific participants were required to be in Canada, the United Kingdom, or the United States, to have an approval rate of greater than 95%, and to be fluent in English.

Participants were asked to perform the experiment in a quiet location and minimize external sounds as much as possible. Next, participants were instructed to set the computer volume to a comfortable level while listening to a calibration noise signal set to the maximum sound level presented during the main experimental task. Each experiment then began with a “headphone check” task to ensure participants were wearing headphones (86). Ensuring headphone use provides more standardized sound presentation across participants and helps to improve overall listening conditions by reducing external background noise. Following the headphone check task, each participant performed a set of practice trials for which feedback was provided after each response. The practice trials helped to ensure participants understood the task instructions and could perform the task correctly. In the main experiment, we incentivized good performance and task engagement by providing feedback after each trial and rewarding participants with a small bonus payment for each correct trial (87).

### Exclusion criteria

During analysis, we screened out participants who were not able to perform the task by excluding those whose performance, averaged across conditions, was below a level that we expected every attentive and normal-hearing participant to achieve, provided they understood the instructions. Because the purpose of the experiments was to assess differences between conditions, rather than absolute performance, this exclusion procedure is neutral with respect to the hypotheses. For Experiments 1 and 4, participants were excluded if average detection performance (d’) was below 0.6. For Experiment 2, participants were excluded if average recognition performance was below 40% correct. For Experiment 3, we planned to exclude participants whose average localization error was above 30° (as it turned out, all had error levels below this criterion). For Experiments 5-10, participants were excluded if average detection performance (d’) was below 0.8. This exclusion criterion excluded between 0% and 17% of participants, depending on the experiment.

### Experiment 1 (Detection)

#### Stimuli

For each of the 160 foreground-background pairs, we constructed stimuli in which the foreground appeared at each of 10 possible temporal positions (foreground onset times of 250, 500, 750, 1000, 1250, 1500, 1750, 2000, 2250, and 2500 ms) and each of 2 possible SNRs (-2 and -6 dB). This yielded a total of 3200 stimuli containing a foreground sound. We also created an additional 160 stimuli that consisted of the background noise only.

#### Procedure

The experiment consisted of 320 trials. Half of these trials included the 160 background noises without a foreground sound. The other half of these trials included each of the 160 foreground-background pairings randomly assigned to one of the 20 experimental conditions (10 foreground positions crossed with 2 SNRs). On each trial, participants judged whether the stimulus contained one or two sound sources.

#### Participants

A total of 200 participants were recruited through Amazon Mechanical Turk. Of these, 88 participants were excluded either because they failed the headphone check task, had self-reported hearing loss, withdrew from the experiment, or completed less than 90% of experimental trials. Finally, 19 participants were excluded due to low task performance (average d’ < 0.6). This resulted in a total of 93 participants included in data analyses. Of these participants, 41 identified as female, 45 as male, and 1 as nonbinary (6 participants did not provide a response). The average age of participants was 37.3 (s.d. = 11.0). All participants were unique to this experiment.

#### Sample size

To determine the sample size necessary to yield stable results, we ran a pilot version of the experiment with 92 participants and calculated the split-half reliability of the average foreground detection performance as we varied the number of participants included in the analysis. The pilot experiment was identical to the actual experiment apart from having 3s background noises (rather than 3.25s as in the actual experiment). Split-half reliability was computed by randomly splitting the sample in half, measuring the Pearson correlation between average performance results in each half, and then applying the Spearman-Brown correction. Because we were primarily interested in the effect of foreground onset time, we measured the split-half reliability separately for each SNR condition then averaged the split-half reliabilities across SNR conditions. Additionally, since the estimated reliability depends on the random split of participants, we repeated this procedure for 10,000 random splits. Because the resulting distribution of reliabilities was skewed, we applied the Fisher z-transform to make the distribution approximately normal. We then took the mean of the Fisher z-transformed distribution (i.e., mean across all random splits) and applied the inverse Fisher z-transformation to obtain our final measure of split-half reliability. We performed this procedure as we varied the number of included participants and found that split-half reliability increased from 0.40 with 10 participants to 0.89 with 92 participants. We fit a curve to these reliabilities and extrapolated that a sample size of 94 participants would be needed to achieve a split-half reliability of at least 0.9. We targeted this sample size, but due to the nature of the screening procedure and the need to collect online data in batches of participants, the actual sample was slightly below this target.

#### Statistics and data analysis

We calculated a hit rate for each of the 20 experimental conditions (10 foreground onset times crossed with 2 SNRs) and a single false alarm rate using the background-only trials. Detection performance was quantified as d’: *d’* = Φ^-1^(Hit Rate) – Φ^-1^(False Alarm Rate) where Φ^-1^ is the inverse CDF of the standard normal distribution. We performed a repeated measures analysis of variance (ANOVA) to analyze the effect of foreground onset time and SNR on foreground detection performance. We assumed data was normally distributed and evaluated this by eye. Mauchly’s test indicated that the assumption of sphericity had not been violated. For each main effect and interaction of interest, we reported F-statistics, p-values and 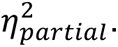 To quantitatively estimate the timescale of improvement with exposure to the background, we fit an elbow function to the results averaged over SNRs. The elbow function was a piecewise linear function consisting of a rise and plateau: 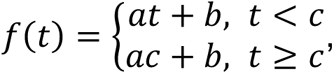 where *a* is the slope of the rise, *b* is the intercept of the rise, *c* is the transition from rise to plateau (i.e., the “elbow point”) and *t* is time. We fit the elbow function by minimizing the absolute error between the estimated elbow function and the data. To obtain a confidence interval around the location of the elbow point, we bootstrapped over participants 10,000 times.

### Experiment 2 (Recognition)

#### Stimuli

We used the same stimuli as in Experiment 1, including only those that contained a foreground sound.

#### Procedure

Each participant heard one trial for each of the 160 foreground-background pairings randomly assigned to one of the 20 experimental conditions (10 foreground positions crossed with 2 SNRs). Thus, the experiment consisted of 160 trials. On each trial, participants were asked to identify the foreground by selecting a text label from five options. One option was the correct label of the foreground, and the remaining options were chosen randomly from the labels of the other foreground sounds in the stimulus set.

#### Participants

A total of 409 participants were recruited through Prolific. Of these, 133 participants were excluded either because they failed the headphone check task, had self-reported hearing loss, withdrew from the experiment, or completed less than 90% of experimental trials. Finally, 15 participants were excluded due to low task performance (average recognition performance < 40% correct). This resulted in a total of 261 participants included in data analyses. Of these participants, 123 identified as female, 134 as male, and 2 as nonbinary (2 participants did not provide a response). The average age of participants was 38.8 (s.d. = 12.1). All participants were unique to this experiment.

#### Sample size

To determine the sample size necessary to yield stable results, we ran a pilot version of the experiment with 103 participants and calculated the split-half reliability of the average foreground recognition performance as we varied the number of participants included in the analysis. The pilot experiment was identical to the actual experiment apart from having 3s background noises (rather than 3.25s as in the actual experiment). The procedure for determining sample size was identical to that of Experiment 1. We found that split-half reliability increased from 0.07 with 10 participants to 0.53 with 102 participants. We fit a curve to these reliabilities and extrapolated that a sample size of 252 participants would be needed to achieve a split-half reliability of at least 0.9. We targeted this sample size, but due to the nature of the screening procedure, the actual sample was slightly above the target sample size.

#### Statistics and data analysis

We performed a repeated measures ANOVA to analyze the effect of foreground onset time and SNR on foreground recognition performance (quantified as precent correct). We assumed data was normally distributed and evaluated this by eye. Mauchly’s test indicated that the assumption of sphericity had not been violated. For each main effect and interaction of interest, we reported F-statistics, p-values and 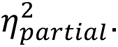 The procedure for fitting the elbow function was identical to that of Experiment 1.

### Experiment 3 (Localization)

#### Stimuli

For each of the 160 background noises, we synthesized five unique 7-second-long exemplars and cut each exemplar into two 3.25-second-long sounds to yield a total of 10 unique waveforms for each background noise. We chose to synthesize 7-s exemplars rather than the 9-s exemplars used to generate stimuli in Experiments 1 and 2 because it reduced the time for synthesis while still enabling two excerpts to be cut from each exemplar. On a given trial, these 10 noise exemplars were played from 10 randomly chosen speakers to create diffuse background noise. Each background noise was played at a level 52 dBA such that the total level of background noise was 62 dBA. The foreground sounds were identical to the 0.5s clips used in previous experiments and were played at a random speaker location (distinct from the 10 locations of the background noise) at a level of 50 dBA (i.e., at an SNR of -12 dB).

#### Procedure

Each participant heard one trial for each of the 160 foreground-background pairings, randomly assigned to one of the five experimental conditions (foreground onset times of 250, 750, 1250, 1750, and 2250 ms). Participants were instructed to fixate on the speaker directly in front of them, with their head still, for the duration of sound presentation. At the end of the sound presentation, participants could move their head to note the label of the speaker from which they judged the foreground sound to have played from. This label was entered using a keyboard. Participants were then instructed to reorient to the speaker directly in front of them before beginning the next trial. Trials were presented in two blocks of 80 trials with a short break between the blocks.

#### Participants

A total of 22 participants were recruited from the area around Cambridge, MA. Of these participants, 7 identified as female and 15 as male. The average age of participants was 26.4 (s.d. = 3.6). All participants were unique to this experiment. All participants provided informed consent and the Massachusetts Institute of Technology Committee on the Use of Humans as Experimental Subjects (COUHES) approved this experiment. No participants were excluded due to low task performance (average localization error > 30°).

#### Sample size

To determine an appropriate sample size, we performed a power analysis using G*Power (88). We sought to be 90% likely to detect an effect as big as that observed in Experiment 1, at a p<0.01 significance level using a repeated measures ANOVA with 5 repeated measurements (foreground onset times), assuming sphericity and a correlation among repeated measures of 0.2 (estimated from Experiment 1). This yielded a target sample size of 17 participants. We ran somewhat more than this to be conservative.

#### Statistics and data analysis

We performed a repeated measures ANOVA to analyze the effect of foreground onset time on foreground localization performance. Localization performance was quantified as the absolute localization error in azimuth. We assumed data was normally distributed and evaluated this by eye. Mauchly’s test indicated that the assumption of sphericity had not been violated. For the main effect of interest, we reported F-statistics, p-values and 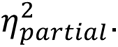 The procedure for fitting the elbow function was identical to that of Experiment 1.

### Experiment 4 (Cued Detection)

#### Stimuli

The stimuli were identical to those of Experiment 1 but with lower SNRs (-5 and -8 dB). The cue sound was always the same waveform as the foreground sound that could appear within the background, the only difference being that the foreground amplitude was scaled to achieve the desired SNR for that trial. The cue was presented at the same level as the background, and thus differed in level from the foreground.

#### Procedure

The experiment consisted of 320 trials. On each trial, participants first heard a foreground sound in isolation (the “cued sound”), followed by continuous background noise. Half of the trials contained the cued foreground sound superimposed somewhere on the background noise, randomly assigned to one of the 20 experimental conditions (10 foreground positions crossed with 2 SNRs). Participants judged whether the stimulus contained the cued sound.

#### Participants

A total of 240 participants were recruited through Amazon Mechanical Turk. Of these, 81 participants were excluded either because they failed the headphone check task, had self-reported hearing loss, withdrew from the experiment, or completed less than 90% of experimental trials. Finally, 23 participants were excluded due to low task performance (average d’ < 0.6). This resulted in a total of 136 participants included in data analyses. Of these participants, 61 identified as female, 68 as male, and 1 as nonbinary (6 participants did not provide a response). The average age of participants was 38.5 (s.d. = 11.3). All participants were unique to this experiment.

#### Sample size

To determine the sample size necessary to yield stable results, we ran a pilot version of the experiment with 95 participants and calculated the split-half reliability of the average foreground detection performance as we varied the number of participants included in the analysis. The pilot experiment was identical to the actual experiment apart from having 3s background noises (rather than 3.25s as in the actual experiment) and SNRs of -2 and -6 dB (rather than -5 and -8 dB in the actual experiment). The procedure for determining sample size was identical to that of Experiment 1. We found that split-half reliability increased from 0.37 with 10 participants to 0.88 with 94 participants. We fit a curve to these reliabilities and extrapolated that a sample size of 105 participants would be needed to achieve a split-half reliability of at least 0.9. We targeted this sample size, but due to the nature of the screening procedure and the need to collect online data in batches of participants, the actual sample was slightly above the target sample size.

#### Statistics and data analysis

We calculated a hit rate for each of the 20 experimental conditions (10 foreground onset times crossed with 2 SNRs) and a single false alarm rate using the background-only trials, then quantified detection performance as d’. We performed a repeated measures ANOVA to analyze the effect of foreground onset time and SNR on foreground detection performance. We assumed data was normally distributed and evaluated this by eye. Mauchly’s test indicated that the assumption of sphericity had not been violated. For each main effect and interaction of interest, we reported F-statistics, p-values and 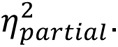 The procedure for fitting the elbow function was identical to that of Experiment 1.

### Observer model

#### Overview

First, an input sound waveform is passed through a standard model of auditory processing consisting of two stages: a peripheral stage modeled after the cochlea, yielding a “cochleagram”, followed by a set of spectrotemporal filters (inspired by the auditory cortex) that operate on the cochleagram, yielding time-varying activations of different spectrotemporal features. Next, a probability distribution is estimated from the filter activations over a past time window. This distribution is then used to evaluate the surprisal of samples in a present time window. The process is then stepped forward in time and repeated, resulting in a set of surprisal values for each time point of the stimulus. Finally, this surprisal curve is compared to a time-varying decision threshold to decide whether a foreground sound is present.

#### Cochleagram

Cochleagrams were computed with a set of 40 filters (38 bandpass filters plus one low-pass and one high-pass filter). Filter cutoffs were evenly spaced on an ERB-scale (89) and thus mirrored the frequency resolution believed to characterize the human cochlea. Filters had transfer functions that were a half-cycle of a cosine function. The cochleagram resulted from the following sequence of steps (24). First, the filters were applied to the audio signal (at an audio sampling rate of 20000 Hz), yielding subbands. Second, subband envelopes were computed using the Hilbert transform. Third, the subband envelopes were passed through a compressive nonlinearity (by raising them to a power of 0.3). Fourth, the compressed envelopes were downsampled to a sampling rate of 2000 Hz.

#### Spectrotemporal filters

We selected spectrotemporal filters that were principal components of a large set of natural textures, as these captured the variance within natural background sounds. We first extracted 100 random 50-ms-long segments from the cochleagram representation of 1000 sound textures not used in our experiments, and then ran principal component analysis on these cochleagram segments. We found that 541 principal components were sufficient to explain 95% of the variance in the random segments and subsequently used these components as the spectrotemporal filters for our model. The filter activations were the dot product of the filter with the stimulus cochleagram.

#### Surprisal

Surprisal is defined as the negative log-probability of an event. Because we model filter activations using a continuous, normal distribution, we calculate surprisal using the negative log-density. For a univariate normal random variable *X* ∼ 𝒩(*μ*, *σ*^2^), surprisal can be written as:

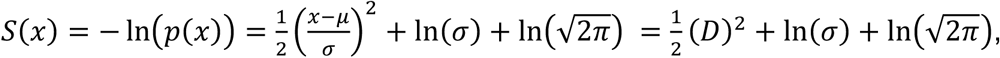

where *D* is the Mahalanobis distance. Thus, any event that occurs with low likelihood will have high surprisal. On the grounds that foreground sounds should be unlikely under a distribution of the background, our model detects the presence of a foreground sound by tracking when the surprisal exceeds some criterion threshold. However, because the surprisal scales with the natural logarithm of the standard deviation, any threshold used for this purpose must similarly scale with the standard deviation of the background. In practice, rather than scale the decision threshold for each stimulus, we instead scale the surprisal by subtracting off the standard deviation term then use a fixed decision threshold across all stimuli.

#### Distribution fitting procedure

Due to the large number of spectrotemporal filters used, fitting a single high-dimensional joint distribution to the activations of all filters was intractable. Thus, we assumed activations across filters to be independent and fit separate univariate distributions to each filter’s activations. In particular, we assumed filter activations were univariate Gaussians. We estimated the mean and variance of the activations within a past window and used these values to calculate the surprisal in a present window (averaging the surprisal over each time point within the window). We repeated this procedure at a sequence of time points, stepping forward in increments of 10 ms. This yielded a surprisal curve (surprisal over time) for each spectrotemporal filter. We then averaged across filters to yield the final surprisal curve. The size of the past window over which distributional parameters are estimated is a model hyperparameter. We tested past window sizes of 500, 750, 1000, 1250, 1500, 1750, 2000, 2250 and 2500 ms and present window sizes of 100, 250 and 500 ms. We found the 1000 ms past window and 500 ms present window to give the best fit with human results, as measured by the correlation with human results. The latter value is intuitively sensible given the 500 ms foreground duration. Figure 3 shows results for these window lengths.

#### Boundary handling

Boundaries pose a challenge for the estimation process in our model (and for the human perceptual system), for two reasons. First, at the onset of a stimulus, there is not yet enough stimulus history with which to estimate distribution parameters for the computation of surprisal (because there are not enough data points to reliably estimate parameters). Second, the filter activations contain boundary artifacts caused by the stages of filtering applied to the stimulus onset. We mitigated these issues by taking a weighted average of the estimated distributional parameters (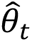 at time *t*) and a prior (*π*_*t*_ at time *t*) whenever the available stimulus history is less than the past window size (*l* in samples) over which parameters are estimated. Because the model was fit to the activations of spectrotemporal filters derived from PCA, the prior on the mean was 0 and the prior on the variance was given by the variance of each principal component across the set of random texture segments from which the principal components were computed (see “Spectrotemporal filters” section above). The weight (*w*_*t*_ at time *t*) was linearly relaxed from full weight on the prior at stimulus onset to full weight on the estimated parameters once the available stimulus history was equal to the size of the past window. Thus, the model parameters (*θ*_*t*_ at time *t*) were given by:

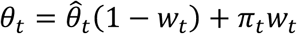

where 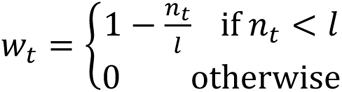 and *n*_*t*_ is the number of samples available at time *t*.

#### Time-varying decision threshold

To determine a decision threshold, we ran the model on 100 random 3.25s excerpts of the 160 textures used in our experiments (see “Simulation of Experiment 1” below). This yielded a total of 16,000 surprisal curves. Then, for each point in time, we took the mean and standard deviation across all surprisal curves to quantify the distribution of surprisal in the absence of a foreground sound. The main idea is that surprisal values greater than that expected by chance (i.e., falling in the tail of this distribution) should indicate the presence of a foreground sound. We thus took the mean plus some number of standard deviations as the decision threshold. The number of standard deviations was chosen via grid search (1000 samples linearly spaced between 0.5 and 5) to best match the model’s false alarm rate to that of human participants in Experiment 1. This was done separately for each set of model hyperparameters (i.e., past and present window sizes).

#### Decision rule

For each point in time, we evaluated whether the measured surprisal exceeded the decision threshold. The model decided a foreground sound was present if the surprisal exceeded the decision threshold for at least 50% of the time in any 500ms window.

#### Simulation of Experiment 1 (Fig. 3B)

Because the model could be run on arbitrarily many stimuli, we opted to show the model results in the limit of a very large amount of data. We simulated the experiment on a larger set of stimuli obtained by generating multiple texture exemplars for each of the background textures used in the human experiments. The stimuli were otherwise identical to those used in Experiment 1. To generate these stimuli, we synthesized 10 unique exemplars of each background noise texture then randomly took 10 different excerpts from each to yield a total of 100 unique excerpts of each background noise. We then ran the model on each of the 3,360 possible stimulus configurations (see “Stimuli” section in Experiment 1 above) for all 100 excerpts of a given background to yield model responses to a total of 336,000 stimuli. To provide a sense of the variability in model results for different subsets of stimuli, we computed model performance (quantified as d’) over 10,000 subsets of the total 336,000 stimuli. Specifically, for each of the 20 experimental conditions, one stimulus was chosen randomly for each of the 160 foreground-background pairings and a model hit rate was computed from these 160 trials. Thus, a total of 3,200 (20 conditions x 160 pairings) trials were used to calculate the model hit rates for each experimental condition. To compute a model false-alarm rate, we randomly selected 20 background-only stimuli for each of the 160 backgrounds, giving another 3,200 trials. Together, this yielded a total of 6,400 stimuli (with half containing a foreground) for which performance was evaluated at each bootstrapped sample. Final model performance was taken as the mean performance across the 10,000 bootstrapped samples.

### Experiment 5a (Short Interruptions in Background Noise)

#### Stimuli

For each of the 160 foreground-background pairs, we constructed 4-second-long stimuli in which the middle 500 ms of background noise was replaced with either silence or white noise (12 dB higher in level relative to the background). The foreground sound appeared at each of 8 possible temporal positions (foreground onset times of 250, 500, 750, 1000, 2500, 2750, 3000, and 3250 ms), at an SNR of -2 dB. This yielded a total of 2560 stimuli containing a foreground sound. We also created an additional 320 stimuli that consisted of the background noise only with each of the two possible “interrupters” (silence or white noise).

#### Procedure

The experiment consisted of 320 trials. Half of these trials presented the 160 background noises without a foreground sound, randomly assigned to one of the two interrupter conditions. The other half of these trials included each of the 160 foreground-background pairings randomly assigned to one of the 16 experimental conditions (8 foreground positions crossed with 2 interrupter types). Participants were instructed to ignore the interrupter and judge whether the stimulus contained one or two sound sources.

#### Participants

A total of 105 participants were recruited through Prolific. Of these, 27 participants were excluded either because they failed the headphone check task, had self-reported hearing loss, withdrew from the experiment, or completed less than 90% of experimental trials. No participants were excluded due to low task performance (average d’ < 0.8). This resulted in a total of 78 participants included in data analyses. Of these participants, 45 identified as female, 32 as male, and 1 as nonbinary. The average age of participants was 38.3 (s.d. = 12.2). All participants were unique to this experiment.

#### Sample size

To determine the sample size necessary to yield stable results, we ran a pilot version of the experiment with 57 participants and calculated the split-half reliability of the average foreground detection performance for foreground onset times prior to the interrupter as we varied the number of participants included in the analysis. The pilot experiment was identical to the actual experiment apart from being run on Mechanical Turk rather than Prolific. At the time the pilot experiment was run, data quality on Mechanical Turk had declined due to an uptick in fraudulent workers, and so we opted to run the actual experiment on Prolific but still considered the Mechanical Turk data to be reasonable as a pilot. The procedure for determining sample size was identical to that of Experiment 1. We found that split-half reliability increased from 0.40 with 10 participants to 0.84 with 56 participants. We fit a curve to these reliabilities and extrapolated that a sample size of 78 participants would be needed to achieve a split-half reliability of at least 0.9.

#### Statistics and data analysis

We calculated a hit rate for each of the 16 experimental conditions (8 foreground onset times crossed with 2 interrupter types) and false alarm rates using the background-only trials for each interrupter type, then quantified detection performance as d’. We performed a repeated measures ANOVA to analyze the effect of interrupter type and foreground position (relative to the interrupter) on foreground detection performance. We assumed data was normally distributed and evaluated this by eye. Mauchly’s test indicated that the assumption of sphericity had not been violated. For each main effect and interaction of interest, we reported F-statistics, p-values and 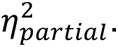

### Experiment 5b (Longer Interruptions in Background Noise)

#### Stimuli

The stimuli were created in a manner similar to that of Experiment 5a. Previous work using EEG to measure adaptation in auditory-evoked cortical potentials in humans found that the recovery from adaptation (in silence) followed an exponential function with a time-constant of around 1300 ms (51). Thus, it seemed possible that the timescale of recovery from adaption exceeded the duration of the 500 ms interrupter used in Experiment 5a, causing the benefit of background exposure to persist across the interruption. Thus, in Experiment 5b, we increased the duration of the interrupter to 1500 ms. For each of the 160 foreground-background pairs, we constructed 5-second-long stimuli in which the middle 1500 ms of background noise was replaced with either silence or white noise (12 dB higher in level relative to the background). Because it seemed plausible that gaps between the noise and the background texture might make the noise more salient, making for a stronger test, the first and last 125 ms of the white noise interrupter was replaced with silence. The foreground sound appeared at each of 8 possible temporal positions (foreground onset times of 250, 500, 750, 1000, 3500, 3750, 4000, and 4250 ms), at an SNR of -2 dB. This yielded a total of 2560 stimuli containing a foreground sound. We also created an additional 320 stimuli that consisted of the background noise only with each of the two possible “interrupters” (silence or white noise).

#### Procedure

The procedure was identical to that of Experiment 5a.

#### Participants

A total of 121 participants were recruited through Prolific. Of these, 49 participants were excluded either because they failed the headphone check task, had self-reported hearing loss, withdrew from the experiment, or completed less than 90% of experimental trials. Finally, 1 participant was excluded due to low task performance (average d’ < 0.8). This resulted in a total of 71 participants included in data analyses. Of these participants, 31 identified as female, 39 as male, and 1 as nonbinary. The average age of participants was 34.3 (s.d. = 10.2). All participants were unique to this experiment.

#### Sample size

Because we planned to compare the results of Experiment 5b to that of Experiment 5a, we targeted the size of the sample collected in Experiment 5a (n=78), but due to the nature of the screening procedure, the actual sample was slightly below the target sample size.

#### Statistics and data analysis

Like Experiment 5a, we calculated a hit rate for each of the 16 experimental conditions (8 foreground onset times crossed with 2 interrupter types) and false alarm rates using the background-only trials for each interrupter type, then quantified detection performance as d’. We performed a repeated measures ANOVA to analyze the effect of interrupter type and foreground position (relative to the interrupter) on foreground detection performance. We assumed data was normally distributed and evaluated this by eye. Mauchly’s test indicated that the assumption of sphericity had not been violated. To compare foreground detection performance following different interrupter durations (between Experiments 5a and 5b), we also performed a mixed model ANOVA with interrupter type as a within-subject factor and interrupter duration as a between-subject factor, including only onset times after the interruption in each experiment. For each main effect and interaction of interest, we reported F-statistics, p-values and 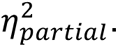

### Experiment 6 (Repeating Background Noises on Every Trial)

#### Stimuli

For each of the 160 background noises, we synthesized 20 7-second-long exemplars and cut each exemplar into two 3.25-second-long sounds to yield a total of 40 unique waveforms for each background noise. For each of these 6,400 unique background noise waveforms, each of the 160 foregrounds could appear at each of 10 possible temporal positions (foreground onset times of 250, 500, 750, 1000, 1250, 1500, 1750, 2000, 2250, and 2500 ms) at an SNR of -8 dB, yielding a total of 10,240,000 possible stimuli containing a foreground sound and 6,400 possible stimuli consisting of background noise only. Rather than create all possible experimental stimuli, we pre-generated enough stimulus sets (see Procedure below) such that each participant in our sample would receive a unique set, generating only the stimuli needed for these sets.

#### Procedure

For each subject, we randomly selected 8 of the 160 possible backgrounds to repeat on every trial in blocks of 40 trials. On half of these trials, the background noise appeared in isolation. The other half of these trials also contained a randomly selected foreground randomly assigned to one of the foreground onset time conditions such that each foreground onset time condition occurred twice during a block. Each background noise was a unique exemplar, and foregrounds were never repeated. The order of the blocks was chosen at random, as was the order of stimuli within a block. On each trial, participants judged whether the stimulus contained one or two sound sources and were not explicitly informed that backgrounds would repeat.

#### Participants

A total of 289 participants were recruited through Prolific. Of these, 93 participants were excluded either because they failed the headphone check task, had self-reported hearing loss, withdrew from the experiment, or completed less than 90% of experimental trials. No participants were excluded due to low task performance (average d’ < 0.8). This resulted in a total of 196 participants included in data analyses. Of these participants, 90 identified as female, 100 as male, and 6 as nonbinary. The average age of participants was 36.5 (s.d. = 11.9). All participants were unique to this experiment.

#### Sample size

We targeted the same sample size as in Experiment 7 (which is presented later in the text, but which was in practice run first), but due to the nature of the screening procedure, the actual sample was slightly below the target sample size.

#### Statistics and data analysis

We calculated a hit rate for each of the 10 experimental conditions (10 foreground onset times) and a single false alarm rate using the background-only trials, then quantified detection performance as d’. We performed a repeated measures ANOVA to analyze the effect of foreground onset time on foreground detection performance. We assumed data was normally distributed and evaluated this by eye. Mauchly’s test indicated that the assumption of sphericity had not been violated. For each main effect and interaction of interest, we reported F-statistics, p-values and 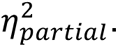

### Experiment 7 (Non-repeated Background Noises with Random Foreground Pairings)

#### Stimuli

For each of the 160 background noises from Experiment 1, we constructed stimuli in which each of the 160 foregrounds appeared at each of 10 possible temporal positions (foreground onset times of 250, 500, 750, 1000, 1250, 1500, 1750, 2000, 2250, and 2500 ms) at an SNR of -8 dB. This yielded a total of 256,000 stimuli containing a foreground sound. We also created an additional 160 stimuli that consisted of the background noise only.

#### Procedure

The experiment consisted of 320 trials. Half of these trials included the 160 background noises without a foreground sound. The other half of these trials contained a randomly selected foreground randomly assigned to one of the foreground onset time conditions. On each trial, participants judged whether the stimulus contained one or two sound sources.

#### Participants

A total of 361 participants were recruited through Prolific. Of these, 158 participants were excluded either because they failed the headphone check task, had self-reported hearing loss, withdrew from the experiment, or completed less than 90% of experimental trials. Finally, 2 participants were excluded due to low task performance (average d’ < 0.8). This resulted in a total of 201 participants included in data analyses. Of these participants, 82 identified as female, 112 as male, and 5 as nonbinary (2 participants did not provide a response). The average age of participants was 37.1 (s.d. = 12.0). All participants were unique to this experiment.

#### Sample size

Because we expected that the randomized foreground-background pairings used in this experiment would increase the variability of the results, we targeted double the sample size of Experiment 1 (n=93) to help ensure sufficient power. Due to the nature of the screening procedure, the actual sample was slightly above the target sample size.

#### Statistics and data analysis

We calculated a hit rate for each of the 10 experimental conditions (10 foreground onset times) and a single false alarm rate using the background-only trials, then quantified detection performance as d’. We performed a repeated measures ANOVA to analyze the effect of foreground onset time on foreground detection performance. We assumed data was normally distributed and evaluated this by eye. Mauchly’s test indicated that the assumption of sphericity had not been violated. To compare foreground detection performance for repeated (Experiment 6) versus non-repeated (Experiment 7) backgrounds, we performed a mixed model ANOVA with foreground onset time as a within-subject factor and background type as a between-subject factor. To compare foreground detection performance for controlled (Experiment 1) versus non-controlled (Experiment 7) foreground-background pairings, we performed a mixed model ANOVA with foreground onset time as a within-subject factor and foreground-background pairing type as a between-subject factor. For each main effect and interaction of interest, we reported F-statistics, p-values and 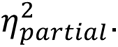 To estimate the overall magnitude of improvement in detection performance with foreground onset time, we fit an elbow function to the results and quantified the delay benefit as the difference between the values of the elbow function at the first (250 ms) and last (2500 ms) foreground onset times. We performed permutation tests to test for differences in the delay benefit across experiments (Experiment 6 versus Experiment 7 or Experiment 1 versus Experiment 7) by randomly shuffling participants across experiments and estimating the difference between the magnitude of improvement in each set of shuffled data. We repeated this procedure 10,000 times to build up a distribution of the test statistic (difference in delay benefit) under the null hypothesis (there is no difference across experiments) and calculated the p-value (two-tailed) as the proportion of times that absolute values from the null distribution were at least as large as the actual absolute difference in delay benefit between experiments. We performed an analogous permutation test to test for a difference in the timescale of improvement (quantified as the location of the elbow point) between Experiments 1 and 7.

To ensure that differences in the delay benefit were not driven by Experiment 6 having more participants with near-ceiling performance compared to Experiment 7, we ran a control analysis in which we selected groups of participants from each experiment to have similar asymptotic performance. To avoid errors of non-independence, we used data from foreground onset times of 1500, 2000, and 2500 ms to select the participant groups, and then measured the delay benefit using the data from the remaining foreground onset times for these participants. In practice, we found that naively matching asymptotic performance for the “selection” conditions (1500, 2000, and 2500 ms) did not result in fully matched performance for the held-out conditions (1250, 1750, and 2250 ms), presumably because the group selection criterion (i.e., the difference in performance between groups for the 1500, 2000, and 2500 ms conditions) had some contribution from noise, which left a residual difference in performance between groups in the held-out conditions. To minimize this difference in performance, we imposed a bias during the matching procedure and selected participant groups whose difference in performance in the selection conditions was as close as possible to the bias value. This bias value was determined by selecting the value which minimized the performance difference between groups via three-fold cross-validation across the three foreground onset times used as the selection conditions. In this way we obtained participant groups whose performance was approximately matched in independent data from the regime in which performance was asymptotic.

### Experiment 8 (Repeating Background Noises on Alternate Trials)

#### Stimuli

The stimuli were identical to those of Experiment 6 but were sampled differently over the course of the experiment due to the constraints of the design (see Procedure below).

#### Procedure

For each subject, we randomly selected 8 of the 160 possible backgrounds to repeat on every other trial in blocks of 40 trials. On half of these trials, the background noise appeared in isolation. The other half of these trials contained a randomly selected foreground randomly assigned to one of the foreground onset time conditions such that each foreground onset time condition occurred once. For the remaining trials, we randomly selected 80 backgrounds to serve as “non-repeating” trials. Each of these backgrounds appeared twice: once in isolation and once with a randomly selected foreground randomly assigned to one of the foreground onset time conditions. These non-repeating background trials were randomly ordered subject to the constraint that each foreground onset time condition (for the non-repeating backgrounds) occurred once during a block. Each background noise was a unique exemplar, and foregrounds were never repeated. The order of the blocks was chosen at random. On each trial, participants judged whether the stimulus contained one or two sound sources and were not explicitly informed that backgrounds would repeat.

#### Participants

A total of 528 participants were recruited through Prolific. Of these, 153 participants were excluded either because they failed the headphone check task, had self-reported hearing loss, withdrew from the experiment, completed less than 90% of experimental trials, or did not complete all critical trials at the start and end of each block. Finally, 7 participants were excluded due to low task performance (average d’ < 0.8). This resulted in a total of 368 participants included in data analyses. Of these participants, 181 identified as female, 178 as male, 5 as nonbinary (4 participants did not provide a response). The average age of participants was 38.3 (s.d.=12.7). All participants were unique to this experiment.

#### Sample size

We targeted the same sample size as in Experiment 6 (n=196) but multiplied it by two to account for the fact that there were half as many trials per condition (20 total conditions: 10 foreground onset times crossed with background type: repeated or non-repeated) in this experiment. Due to the nature of the screening procedure, the actual sample was slightly below the target sample size.

#### Statistics and data analysis

We calculated a hit rate for each of the 20 experimental conditions (10 foreground onset times crossed with 2 background types) and false alarm rates using the background-only trials for each background type (repeated or non-repeated), then quantified detection performance as d’. We performed a repeated measures ANOVA to analyze the effects of foreground onset time and background repetition on foreground detection performance. We assumed data was normally distributed and evaluated this by eye. Mauchly’s test indicated that the assumption of sphericity had not been violated. For each main effect and interaction of interest, we reported F-statistics, p-values and 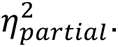 To test for a difference in the delay benefit between background types (repeated versus non-repeated), we performed a permutation test using the same procedure described above in Experiment 7. The control analysis matching asymptotic performance was also performed using the same procedure described above for Experiment 7. However, we note that performing this analysis for Experiment 8 necessitated using distinct but partially overlapping sets of participants for the repeated and non-repeated conditions, because the same participants completed both conditions in the original experiment.

### Experiment 9 (Stationary Noise)

#### Stimuli

To create stationary noise backgrounds, we replaced each of the 160 texture backgrounds with spectrally matched noise. The spectrally matched noise was generated by setting the Fourier amplitudes of a noise signal equal to the Fourier amplitudes of the corresponding sound texture, randomizing the phases, and then performing the inverse Fourier transform. For each of the 160 foreground-background pairs, we constructed stimuli in which the foreground appeared at each of 10 possible temporal positions (foreground onset times of 250, 500, 750, 1000, 1250, 1500, 1750, 2000, 2250, and 2500 ms) and each of 2 possible SNRs (-6 and -10 dB). This yielded a total of 3200 stimuli containing a foreground sound. We also created an additional 160 stimuli that consisted of the stationary background noise only.

#### Procedure

The procedure was identical to that of Experiment 1.

#### Participants

A total of 294 participants were recruited through Prolific. Of these, 86 participants were excluded either because they failed the headphone check task, had self-reported hearing loss, withdrew from the experiment, or completed less than 90% of experimental trials. Finally, 3 participants were excluded due to low task performance (average d’ < 0.8). This resulted in a total of 205 participants included in data analyses. Of these participants, 84 identified as female, 119 as male, and 2 as nonbinary. The average age of participants was 37.6 (s.d. = 12.3). All participants were unique to this experiment.

#### Sample size

We did not have pilot data for this experiment. Thus, we targeted double the sample size of Experiment 1 because of the possibility that the effect of foreground onset time might be smaller. Due to the nature of the screening procedure, we were slightly above the target sample size.

#### Statistics and data analysis

We calculated a hit rate for each of the 20 experimental conditions (10 foreground onset times crossed with 2 SNRs) and a single false alarm rate using the background-only trials, then quantified detection performance as d’. We performed a repeated measures ANOVA to analyze the effect of foreground onset time and SNR on foreground detection performance. We assumed data was normally distributed and evaluated this by eye. Mauchly’s test indicated that the assumption of sphericity had not been violated. For each main effect and interaction of interest, we reported F-statistics, p-values and 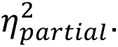 To analyze the effect of stationarity on the pattern of foreground detection performance, we compared the results of Experiment 9 (more stationary spectrally matched noise backgrounds) to the pooled results of Experiments 1 and 7 (less stationary texture backgrounds; data from Experiments 1 and 7 were pooled to increase power given that they showed very similar results). To test for differences in both the magnitude and the timescale of improvement in foreground detection performance between background types (textures versus spectrally matched noises), we performed permutation tests using the same procedure described above in Experiment 7. The control analysis matching asymptotic performance was performed using the same procedure described above in Experiment 7.

### Foreground-Background Similarity Analysis

#### Selection of stimulus pairings

We divided the foreground-background pairings from Experiment 7 (in which foregrounds and backgrounds were randomly paired) into two groups. The groups were selected to be matched in a measure of spectral difference between foreground and background, but to differ in the difference between foreground and background in a spectrotemporal filter basis. Specifically, for each foreground and background, we measured the mean excitation pattern as a summary measure of the spectrum. We also measured the power in each of the 541 PCA-derived spectrotemporal filters used in the observer model. Then, for each foreground-background pair, we computed the Euclidean distance between the excitation patterns and between the spectrotemporal filter powers for the two sounds. Because the two distances are on different scales, we performed min-max normalization for each to scale them between 0 and 1. Next, to select pairings, we calculated the ratio of the spectrotemporal distance to the spectral distance. This ratio is largest when the spectrotemporal distance is large and the spectral distance is small. We then used this measure to split pairings into two groups (low and high spectrotemporal similarity) subject to the constraint that the two groups of pairings contained the same number of occurrences of each foreground and background. This ensures that the results we see are due to the pairings rather than to differences in the specific foregrounds or backgrounds in the two stimulus groups. The free parameter in this procedure was the proportion of total pairings included in the two groups (including all pairings maximized the number of pairings in each group, but led to a smaller difference between groups than if not all pairings were included. We opted to use 75% of all possible pairings, discarding the middle 25% of pairings surrounding the median spectrotemporal distance. This yielded groups containing 9,600 pairings. Because participants were presented with randomly chosen pairings, not all of these 9,600 pairings had been presented to participants (6,923 of the large spectrotemporal distance group and 6,854 of the small spectrotemporal distance group were actually used in the experiment). Figure S5 plots the results separately for trials whose stimuli fell into one group or the other.

#### Statistics and data analysis

Re-analyzing the data from Experiment 7, we calculated a hit rate for each of the 2 groups of pairings for each of the 10 foreground onset times. Using the false alarm rate from the background-only trials of Experiment 7, we quantified detection performance as d’. We performed a repeated measures ANOVA to analyze the effect of foreground onset time and foreground-background similarity on foreground detection performance. We assumed data was normally distributed and evaluated this by eye. Mauchly’s test indicated that the assumption of sphericity had not been violated. For each main effect and interaction of interest, we reported F-statistics, p-values and 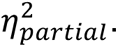 To test for differences in the timescale of improvement in foreground detection performance between pairing types (more similar versus less similar), we performed permutation tests using the same procedure described above for Experiment 7.

### Experiment 10 (Harmonic Foregrounds)

#### Stimuli

To obtain a set of (approximately) harmonic foreground sounds, we selected human vocalizations and musical instrument sounds from a dataset of isolated sound events (GISE-51 training set (90)). Specifically, we selected human vocalization sounds from the “human_speech”, “laughter” and “screaming” categories and selected musical instrument sounds from the “gong”, “guitar”, “harmonica”, “harp”, “marimba_and_xylophone”, “organ”, “piano”, and “trumpet” categories. For each sound in the training set category, we used YIN (54) to measure the average periodicity (one minus aperiodicity) in a sliding 0.5 s window, discarding windowed segments that were mostly quiet or were outside of a periodicity range of 0.9 to 0.99. We set the lower bound to ensure selected sounds would be highly periodic and set the upper bound because windowed segments whose periodicity was greater than 0.99 tended to be tones (e.g., dial tones present in clips labeled as “human_speech”) rather than speech or musical instruments. From the windowed segments of a given sound, we selected the segment with the maximum periodicity (i.e., the most harmonic segment). This resulted in a single 0.5 s clip for each sound in the training set categories described above. Finally, we removed any sounds that were near duplicates (by measuring the power in a set of spectrotemporal filters, computing the correlation of spectrotemporal filter power across sounds, and removing sounds whose correlation exceeded 0.8) or had an estimated F0 below 100 Hz or above 1000 Hz. From the sounds that remained, we chose the most periodic from each category, selecting 40 examples of human speech, 20 examples of laughter and screaming each, as well as 10 examples from each musical instrument category (see Table S3 for a list of the selected sounds).

#### Procedure

The procedure was identical to that of Experiment 7.

#### Participants

A total of 485 participants were recruited through Prolific. Of these, 193 participants were excluded either because they failed the headphone check task, had self-reported hearing loss, withdrew from the experiment, or completed less than 90% of experimental trials. Finally, 5 participants were excluded due to low task performance (average d’ < 0.8). This resulted in a total of 287 participants included in data analyses. Of these participants, 127 identified as female, 151 as male, and 3 as nonbinary (6 participants did not provide a response). The average age of participants was 32.2 (s.d. = 9.7). All participants were unique to this experiment.

#### Sample size

Because we did not have pilot data for this experiment, we initially targeted a sample size double that of Experiment 1 to account for the possibility that the effect of foreground onset time might be smaller. However, we found it difficult to reliably fit elbow functions to this data because the effect of foreground onset time was so small. Thus, we increased the sample size by about 50% to improve the reliability of the elbow function fits to this data.

#### Statistics and data analysis

We calculated a hit rate for each of the 10 experimental conditions (10 foreground onset times) and a single false alarm rate using the background-only trials, then quantified detection performance as d’. We performed a repeated measures ANOVA to analyze the effect of foreground onset time on foreground detection performance. We assumed data was normally distributed and evaluated this by eye. Mauchly’s test indicated that the assumption of sphericity had not been violated. For each main effect and interaction of interest, we reported F-statistics, p-values and 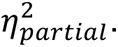 To analyze the effect of harmonicity on the pattern of foreground detection performance, we compared the results of Experiment 10 (harmonic foregrounds) to the pooled results of Experiments 1 and 7 (less harmonic foregrounds; data from Experiments 1 and 7 were pooled to increase power given that they showed very similar results). To test for differences in both the magnitude and the timescale of improvement in foreground detection performance between foreground types, we performed permutation tests using the same procedure described above in Experiment 7. The control analysis matching asymptotic performance was also performed using the same procedure described above in Experiment 7.

## Acknowledgments

We thank the members of the laboratory of J.H.M. for helpful comments on the manuscript, and Andrew Francl, Preston Hess, and Ajani Stewart for building the speaker array used in Experiment 3. This work was supported by NSF grant BCS-2240406 and an NSF Graduate Research Fellowship to J.M.H.

## SI Figures & Tables

**Figure S1.**
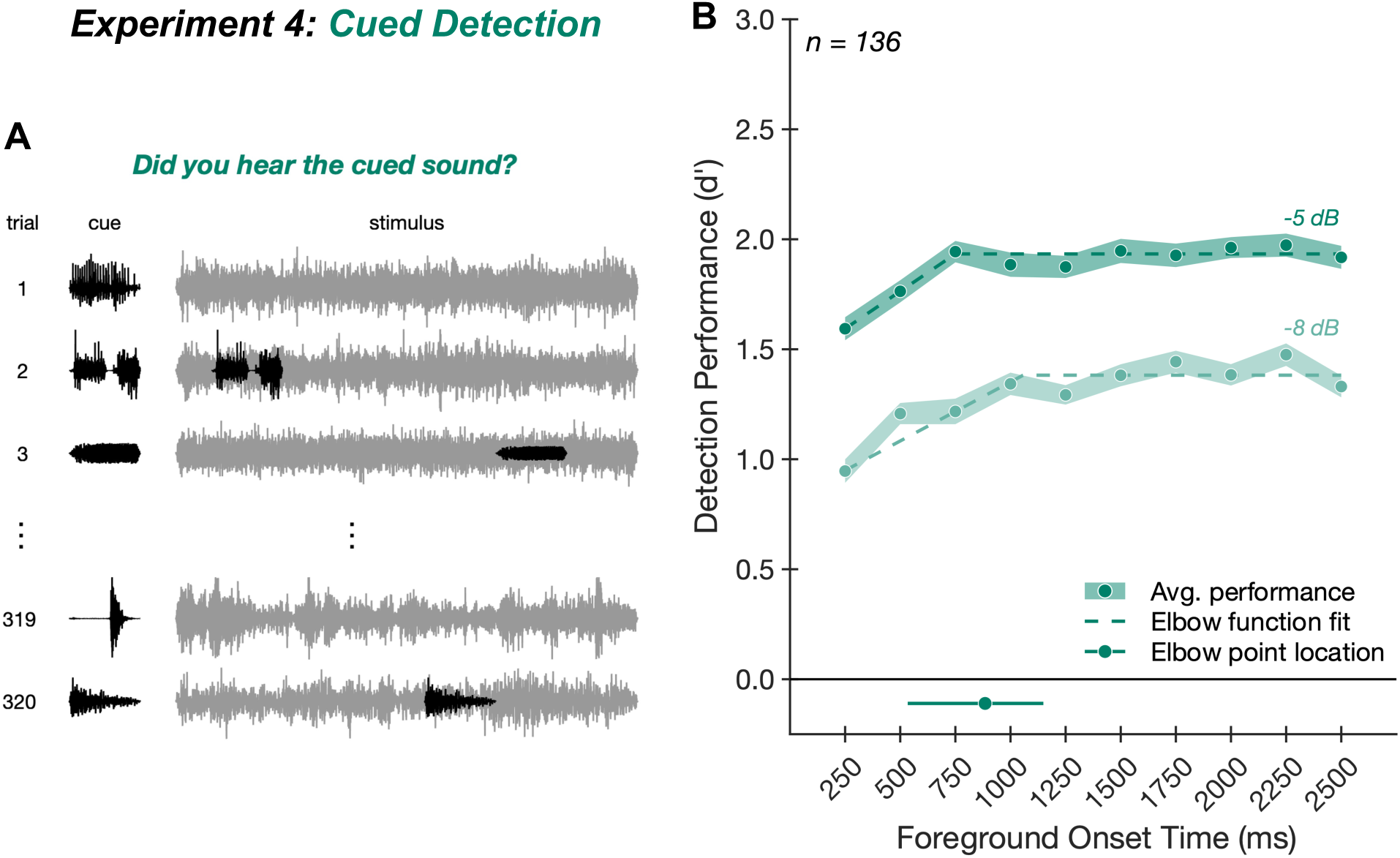
Experiment 4: Benefit of background exposure persists despite knowing what to listen for. (*A*) Experiment 4 task. On each trial, participants first heard a foreground sound in isolation (left, black), followed by continuous background noise (right, gray). Half of the trials contained the cued sound superimposed on the background (e.g., trial 2), and participants judged whether the stimulus contained the cued sound. Because detection performance typically benefits from knowing what to listen for (91), we reduced the SNR of the foreground relative to the background to approximately match the level of performance observed in Experiment 1. (*B*) Experiment 4 results. Average foreground detection performance (quantified as d’; green circles) is plotted as a function of SNR and foreground onset time. Shaded regions plot standard errors. Dashed lines plot elbow function fit. Solid line below main axis plots one standard deviation above and below the median elbow point, obtained by fitting elbow functions to the results averaged over SNR and bootstrapping over participants; dot on this line plots the fitted elbow point from the complete participant sample.

**Figure S2.**
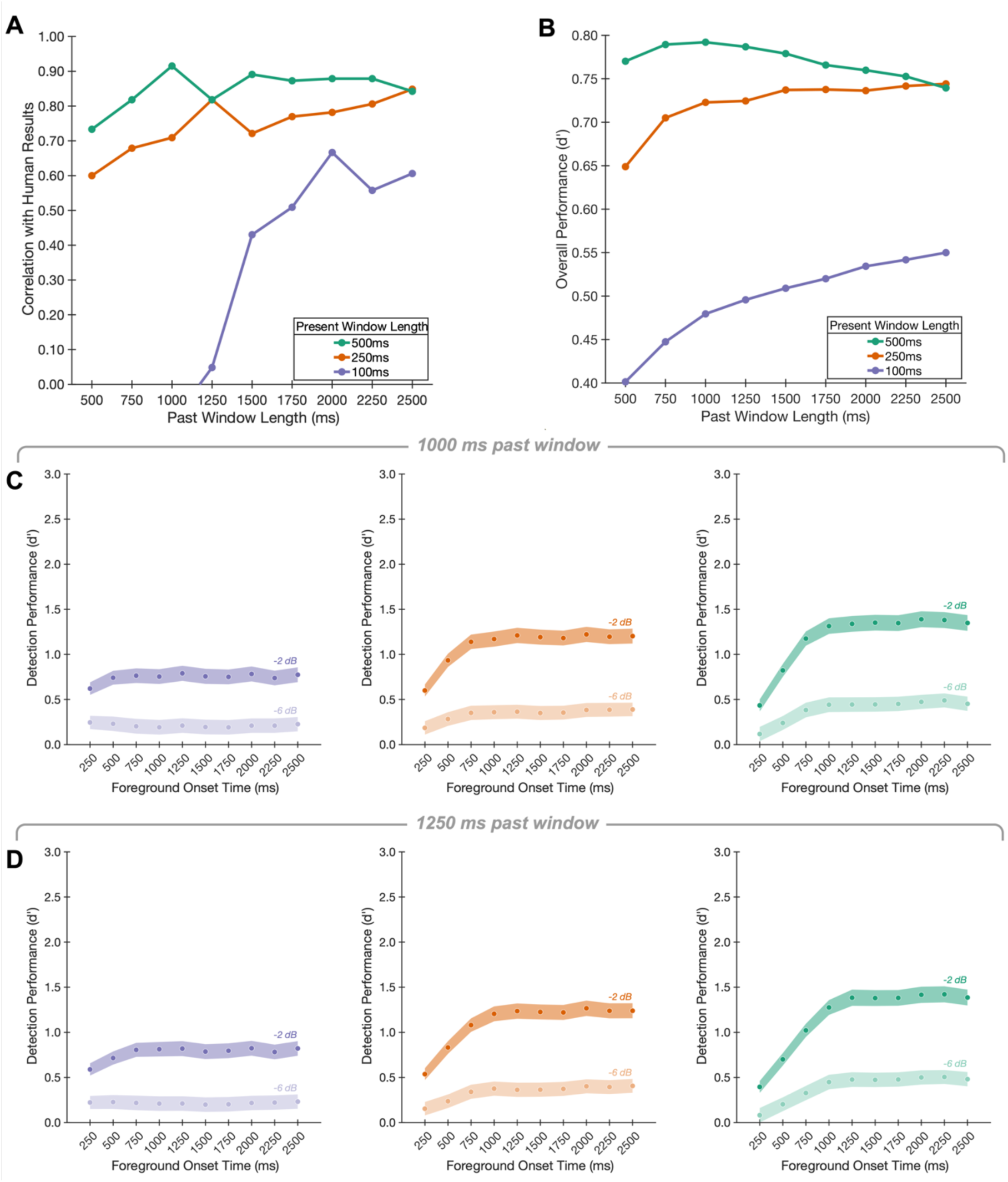
Observer model results for different window sizes. (*A*) Human-model correlations for different window sizes. The Spearman correlation between the model results and the human results from Experiment 1 is plotted as a function of past window length for each present window length (100ms in purple, 250 ms in orange, and 500 ms in green). (*B*) Overall model performance for different window sizes. The overall model performance (computed by averaging detection performance over SNR and foreground onset time) is plotted as a function of past window length for each present window length. Same conventions as *A*. (*C*) Model results using 1000 ms past window. Each panel plots model foreground detection performance as a function of SNR and foreground onset time for a different present window length (100ms left, purple; 250 ms middle, orange; 500 ms right, green) using a fixed past window length of 1000 ms. Shaded regions plot standard deviations of performance obtained by bootstrapping over stimuli. (*D*) Model results using 1250 ms past window. Same conventions as *C* but using a fixed past window length of 1250 ms.

**Figure S3.**
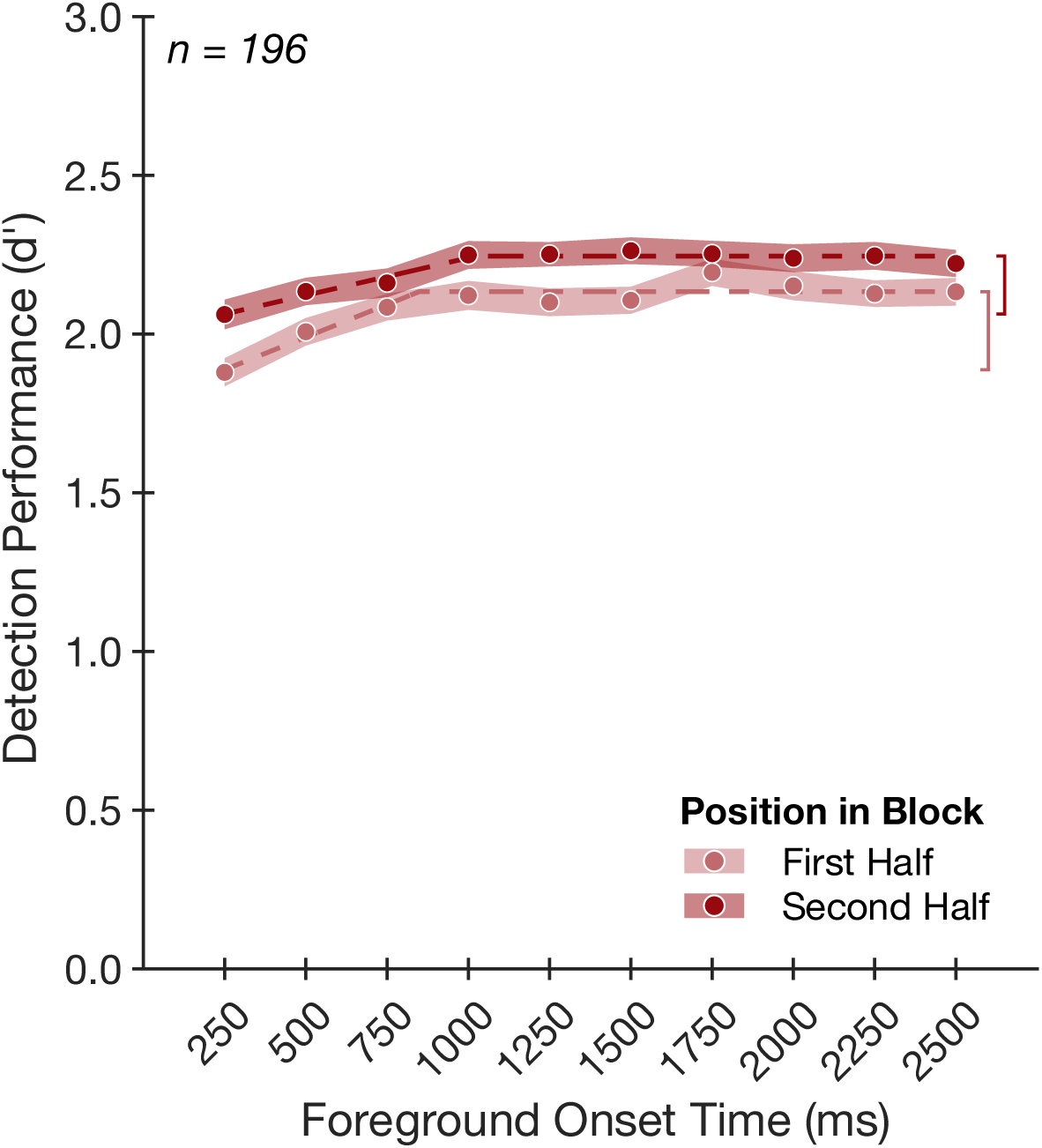
Experiment 6: Repetition of background noise enhances foreground detection. Average foreground detection performance (quantified as d’) is plotted as a function of foreground onset time for the first half of trials (light red circles) versus the second half of trials (dark red circles) within a block. Shaded regions plot standard errors. Dashed lines plot elbow function fit. Vertical brackets denote the delay benefit.

**Figure S4.**
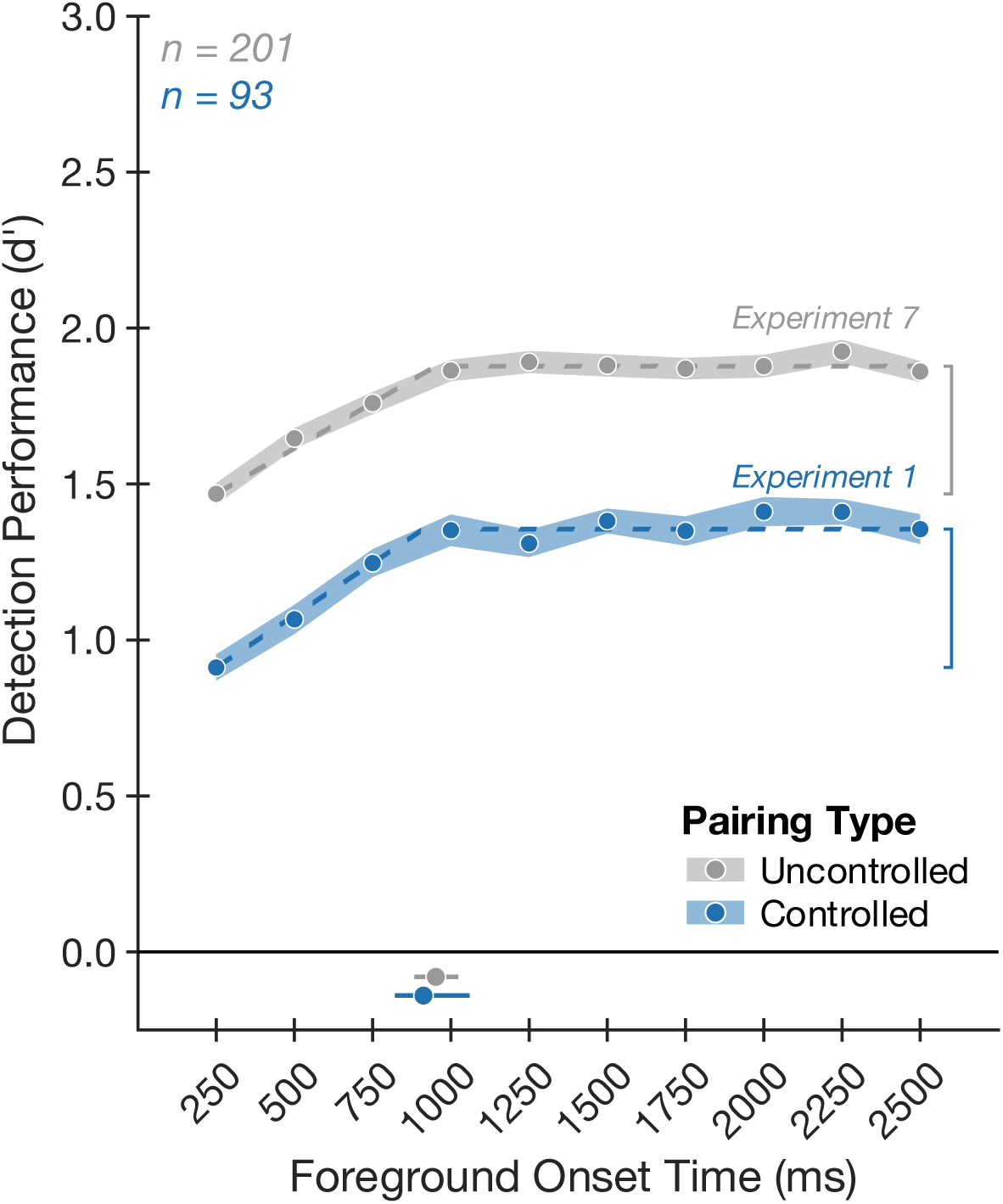
Experiments 1 & 7: Benefit of exposure to background noise is unaffected by choice of foreground-background pairings. Average foreground detection performance (quantified as d’) is plotted as a function of foreground onset time for Experiment 1 (controlled foreground-background pairings; blue circles) versus Experiment 7 (uncontrolled foreground-background pairings; gray circles). Shaded regions plot standard errors. Dashed lines plot elbow function fit. Solid lines below main axis plot one standard deviation above and below the median elbow points (Experiment 1 shown in blue; Experiment 7 shown in gray), obtained by fitting elbow functions to the results averaged over SNR and bootstrapping over participants; dots on these lines plot the fitted elbow points from the complete participant samples. Vertical brackets denote the delay benefit.

**Figure S5.**
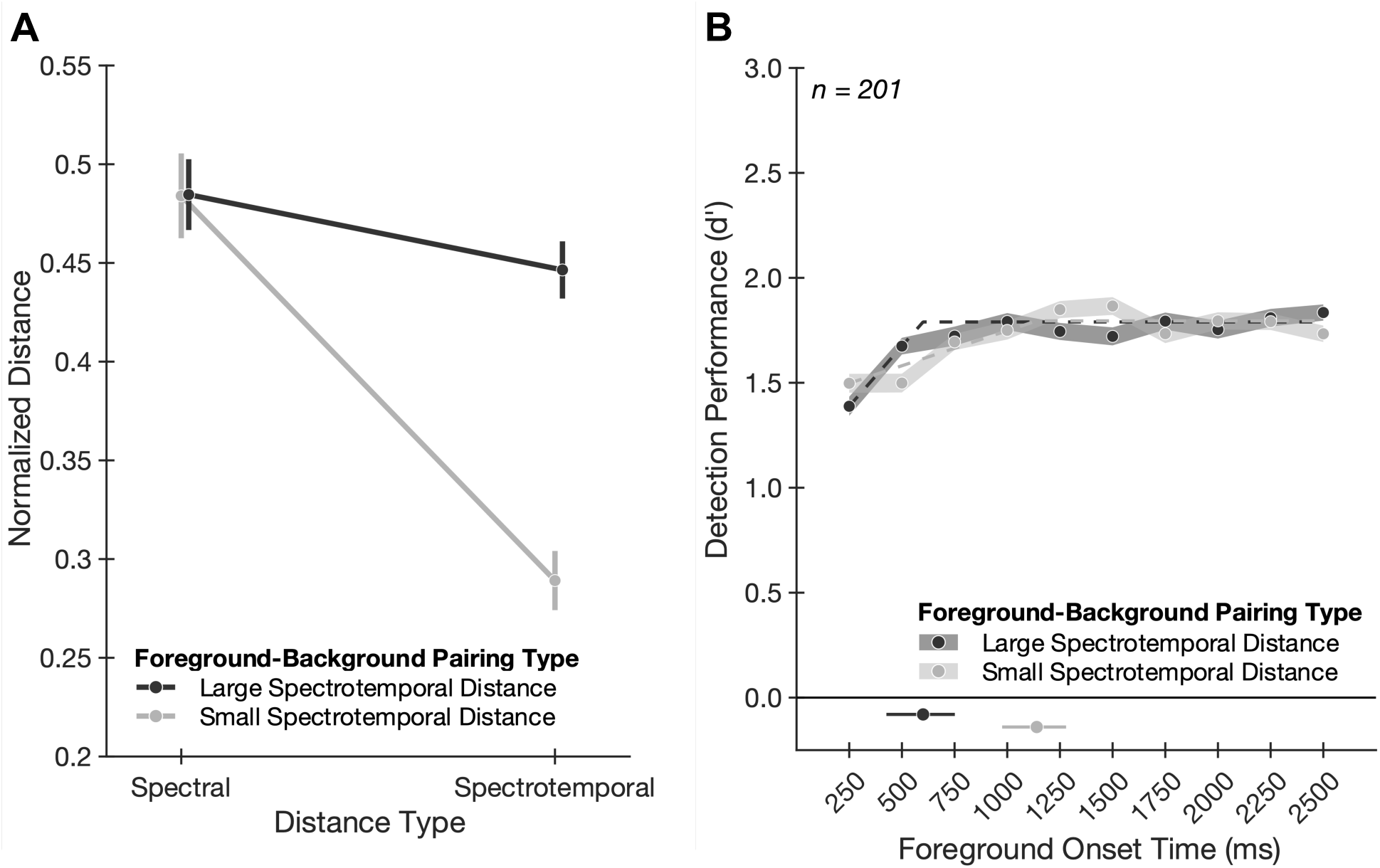
Effect of background exposure depends on foreground-background similarity. (*A*) Spectral and spectrotemporal similarity of foreground-background pairs. The average normalized distance between foregrounds and backgrounds is plotted as a function of distance type for two groups of foreground-background pairings. The two groups of pairings were approximately matched in spectral distance but differed in spectrotemporal distance. Error bars plot standard deviations. (*B*) Foreground-background similarity results. Average foreground detection performance (quantified as d’) is plotted as a function of foreground onset time for the two groups of foreground-background pairs. Shaded regions plot standard errors. Dashed lines plot elbow function fit. Solid lines below main axis plot one standard deviation above and below the median elbow points (more similar pairs shown in gray; less similar pairs shown in black), obtained by bootstrapping over participants; dots on these lines plot the fitted elbow points from the complete participant samples.

**Table S1.**
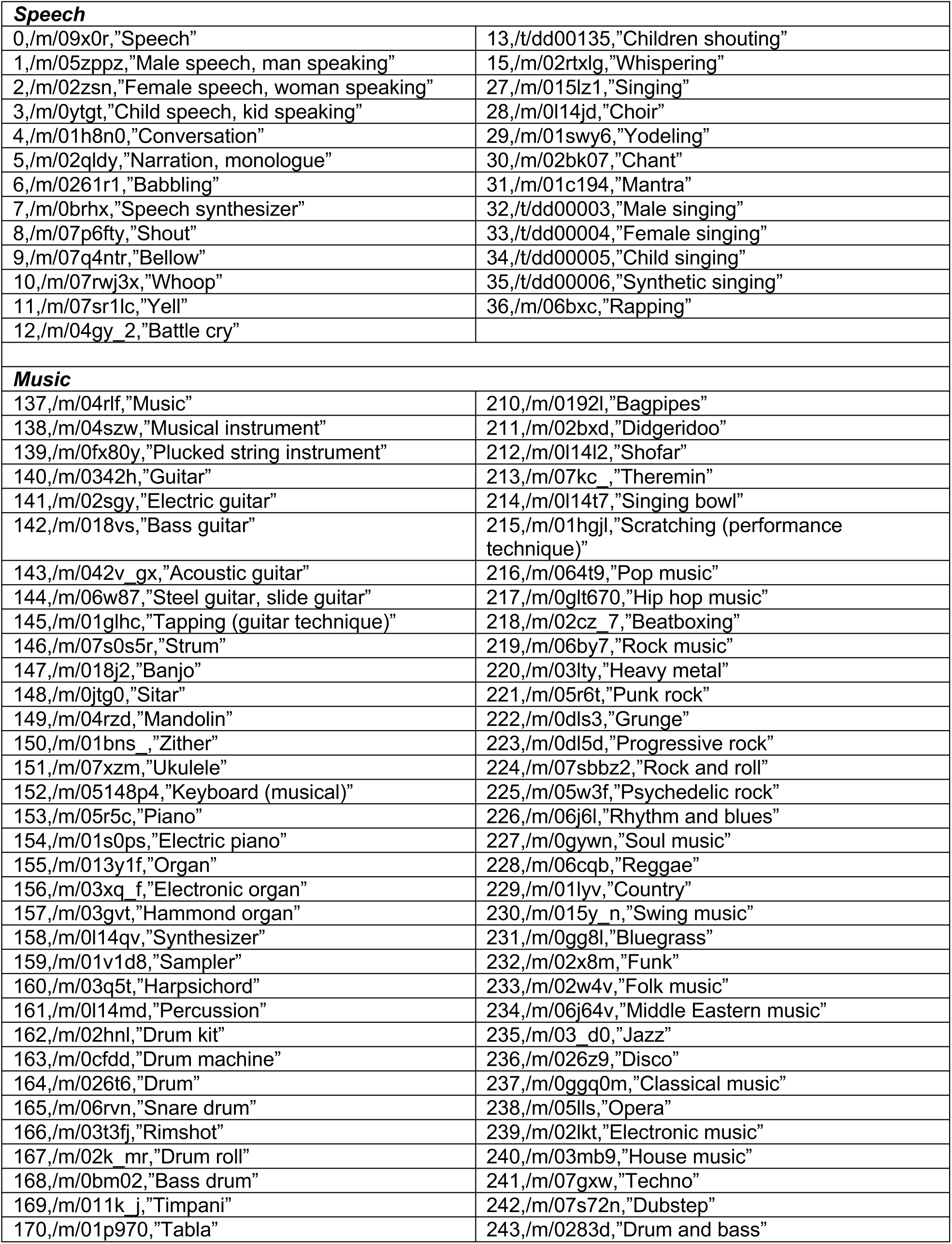
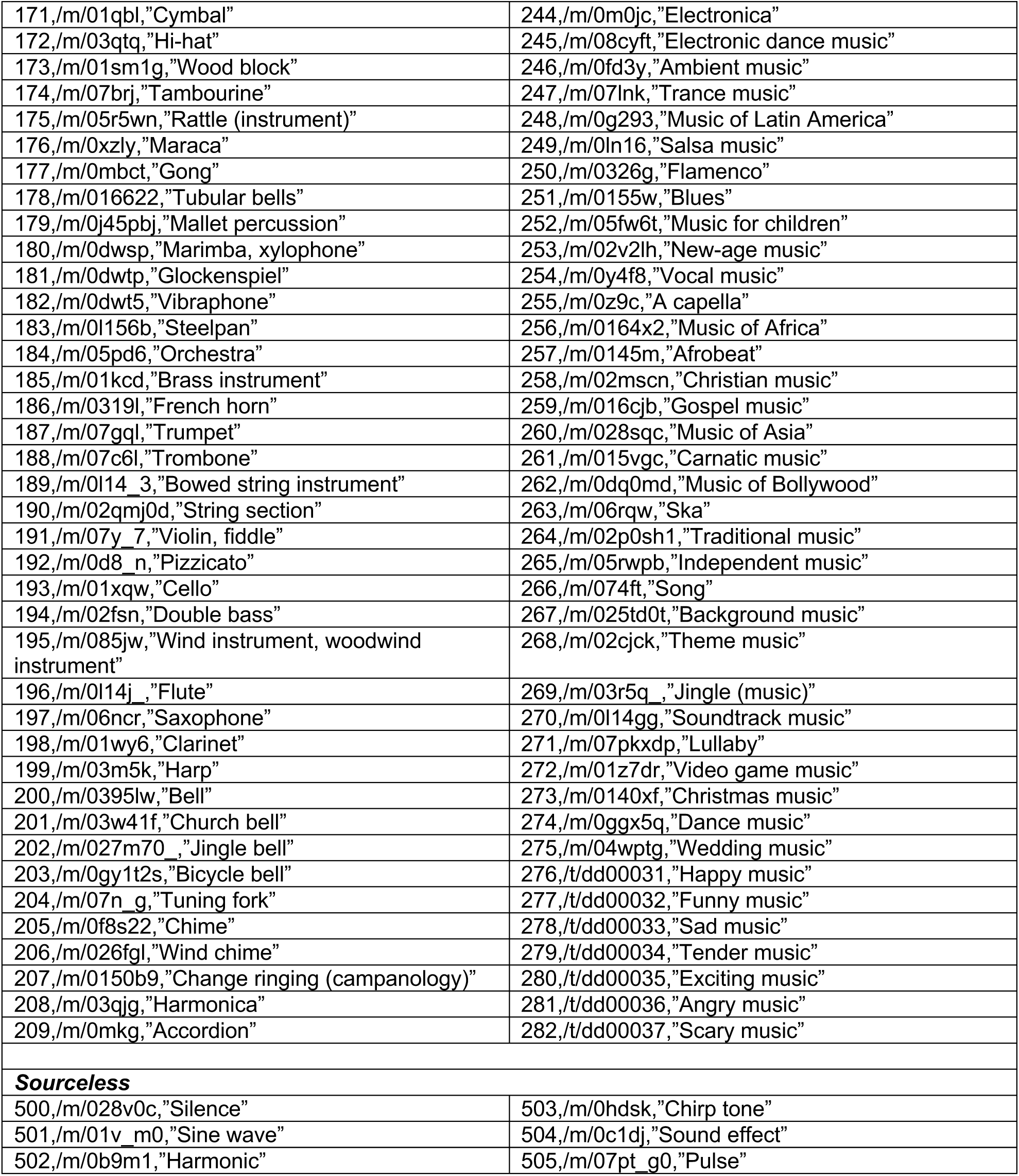
AudioSet labels that were excluded in the process of obtaining texture sounds from which the background noises were drawn.

**Table S2.**
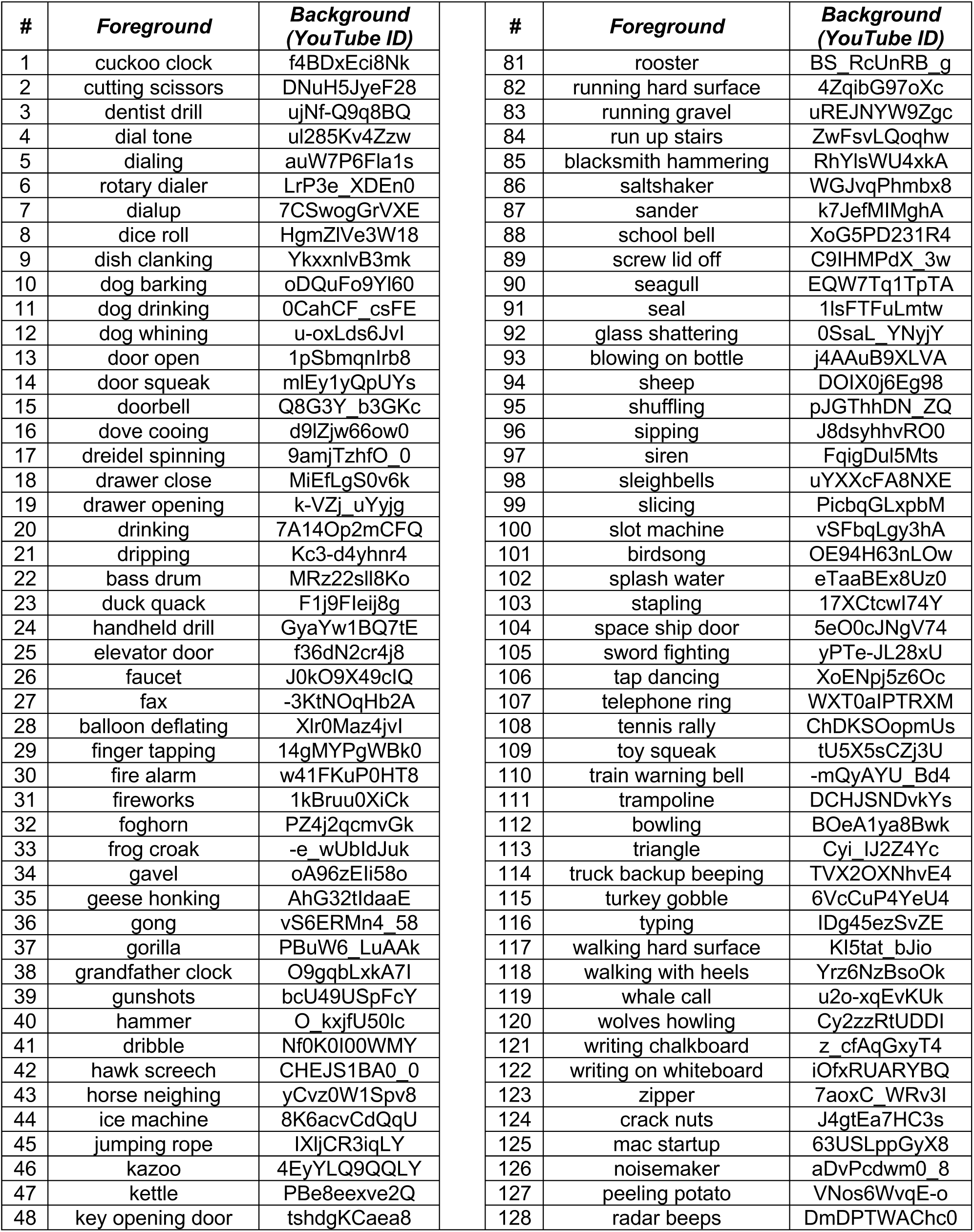
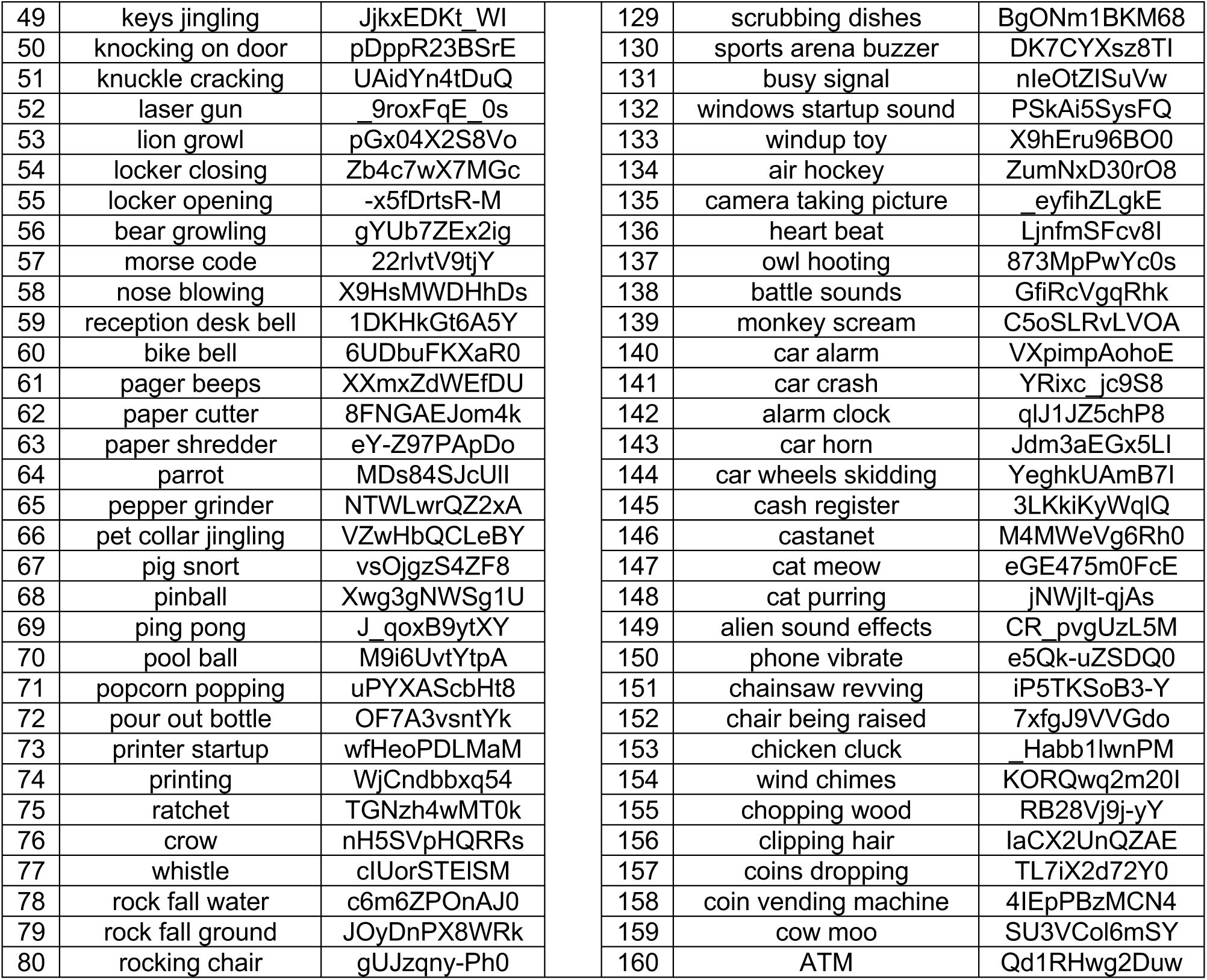
Foreground-background pairings used in Experiments 1-5 and 9. Experiments 6-8 used the same set of foregrounds and backgrounds but paired randomly. Experiment 10 used the same backgrounds but different foregrounds (see Table S3).

**Table S3.** Foregrounds used in Experiment 10, taken from GISE-51 training set.

| # | Category | Stimulus ID | # | Category | Stimulus ID |
| --- | --- | --- | --- | --- | --- |
| 1 | Human speech | 107100 | 81 | Gong | 209917_filtered |
| 2 | Human speech | 92940 | 82 | Gong | 222192 |
| 3 | Human speech | 80641 | 83 | Gong | 57389 |
| 4 | Human speech | 63334 | 84 | Gong | 222191 |
| 5 | Human speech | 181948_filtered | 85 | Gong | 57419 |
| 6 | Human speech | 106439 | 86 | Gong | 206023 |
| 7 | Human speech | 32168 | 87 | Gong | 57424 |
| 8 | Human speech | 93787_filtered | 88 | Gong | 222923_filtered |
| 9 | Human speech | 91076 | 89 | Gong | 222217 |
| 10 | Human speech | 278944 | 90 | Gong | 222922_filtered |
| 11 | Human speech | 95238 | 91 | Guitar | 128844_filtered |
| 12 | Human speech | 77409 | 92 | Guitar | 40438 |
| 13 | Human speech | 109838_filtered | 93 | Guitar | 251187_filtered |
| 14 | Human speech | 82710 | 94 | Guitar | 64791_filtered |
| 15 | Human speech | 50497_filtered | 95 | Guitar | 251161_filtered |
| 16 | Human speech | 216567 | 96 | Guitar | 28399 |
| 17 | Human speech | 108875 | 97 | Guitar | 251156_filtered |
| 18 | Human speech | 92951 | 98 | Guitar | 48355 |
| 19 | Human speech | 92961 | 99 | Guitar | 252448 |
| 20 | Human speech | 18287 | 100 | Guitar | 18474 |
| 21 | Human speech | 77416 | 101 | Harmonica | 116903_filtered |
| 22 | Human speech | 235110 | 102 | Harmonica | 116897_filtered |
| 23 | Human speech | 77421 | 103 | Harmonica | 116902 |
| 24 | Human speech | 184432_filtered | 104 | Harmonica | 325376 |
| 25 | Human speech | 92941 | 105 | Harmonica | 330759_filtered |
| 26 | Human speech | 371581 | 106 | Harmonica | 116887_filtered |
| 27 | Human speech | 235114 | 107 | Harmonica | 325377 |
| 28 | Human speech | 422718_filtered | 108 | Harmonica | 116874_filtered |
| 29 | Human speech | 104707 | 109 | Harmonica | 116875_filtered |
| 30 | Human speech | 92965 | 110 | Harmonica | 116869 |
| 31 | Human speech | 109865 | 111 | Harp | 53854 |
| 32 | Human speech | 104728 | 112 | Harp | 53867 |
| 33 | Human speech | 107099 | 113 | Harp | 53846 |
| 34 | Human speech | 77413 | 114 | Harp | 53864 |
| 35 | Human speech | 67627 | 115 | Harp | 373563_filtered |
| 36 | Human speech | 106428 | 116 | Harp | 53866 |
| 37 | Human speech | 181316 | 117 | Harp | 415387 |
| 38 | Human speech | 106440 | 118 | Harp | 415463 |
| 39 | Human speech | 77412 | 119 | Harp | 415433 |
| 40 | Human speech | 92919 | 120 | Harp | 415466 |
| 41 | Laughter | 132812 | 121 | Marimba_and_xylophone | 373585_filtered |
| 42 | Laughter | 364485_filtered | 122 | Marimba_and_xylophone | 202584 |
| 43 | Laughter | 171707 | 123 | Marimba_and_xylophone | 216753_filtered |
| 44 | Laughter | 95814 | 124 | Marimba_and_xylophone | 202576 |
| 45 | Laughter | 174625 | 125 | Marimba_and_xylophone | 216755_filtered |
| 46 | Laughter | 411082 | 126 | Marimba_and_xylophone | 216761_filtered |
| 47 | Laughter | 198263 | 127 | Marimba_and_xylophone | 216752_filtered |
| 48 | Laughter | 266004 | 128 | Marimba_and_xylophone | 202577 |
| 49 | Laughter | 57734 | 129 | Marimba_and_xylophone | 216751_filtered |
| 50 | Laughter | 172923_filtered | 130 | Marimba_and_xylophone | 216756_filtered |
| 51 | Laughter | 132809_filtered | 131 | Organ | 245073 |
| 52 | Laughter | 364444_filtered | 132 | Organ | 373682_filtered |
| 53 | Laughter | 95816 | 133 | Organ | 160297 |
| 54 | Laughter | 62257_filtered | 134 | Organ | 11856 |
| 55 | Laughter | 198252_filtered | 135 | Organ | 373685_filtered |
| 56 | Laughter | 132746_filtered | 136 | Organ | 373688 |
| 57 | Laughter | 393342_filtered | 137 | Organ | 160301 |
| 58 | Laughter | 366173 | 138 | Organ | 373691_filtered |
| 59 | Laughter | 196066 | 139 | Organ | 373687_filtered |
| 60 | Laughter | 9557 | 140 | Organ | 333753 |
| 61 | Screaming | 219665 | 141 | Piano | 148539_filtered |
| 62 | Screaming | 163729_filtered | 142 | Piano | 11655_filtered |
| 63 | Screaming | 220665 | 143 | Piano | 173763_filtered |
| 64 | Screaming | 219663 | 144 | Piano | 351928 |
| 65 | Screaming | 59163 | 145 | Piano | 11656_filtered |
| 66 | Screaming | 131709 | 146 | Piano | 382533_filtered |
| 67 | Screaming | 219666_filtered | 147 | Piano | 32019_filtered |
| 68 | Screaming | 9434_filtered | 148 | Piano | 11644_filtered |
| 69 | Screaming | 58794 | 149 | Piano | 83132 |
| 70 | Screaming | 104035 | 150 | Piano | 11626_filtered |
| 71 | Screaming | 222586 | 151 | Trumpet | 374138_filtered |
| 72 | Screaming | 351630 | 152 | Trumpet | 357321_filtered |
| 73 | Screaming | 104028_filtered | 153 | Trumpet | 357560_filtered |
| 74 | Screaming | 261853_filtered | 154 | Trumpet | 247248_filtered |
| 75 | Screaming | 166154_filtered | 155 | Trumpet | 247220_filtered |
| 76 | Screaming | 42849_filtered | 156 | Trumpet | 247276_filtered |
| 77 | Screaming | 221151 | 157 | Trumpet | 357419 |
| 78 | Screaming | 104033 | 158 | Trumpet | 247216_filtered |
| 79 | Screaming | 220291 | 159 | Trumpet | 247110 |
| 80 | Screaming | 61219 | 160 | Trumpet | 247381 |

